# Multi-tissue analysis reveals short tandem repeats as ubiquitous regulators of gene expression and complex traits

**DOI:** 10.1101/495226

**Authors:** Stephanie Feupe Fotsing, Jonathan Margoliash, Catherine Wang, Shubham Saini, Richard Yanicky, Sharona Shleizer-Burko, Alon Goren, Melissa Gymrek

## Abstract

Short tandem repeats (STRs) have been implicated in a variety of complex traits in humans. However, genome-wide studies of the effects of STRs on gene expression thus far have had limited power to detect associations and provide insights into putative mechanisms. Here, we leverage whole genome sequencing and expression data for 17 tissues from the Genotype-Tissue Expression Project (GTEx) to identify STRs for which repeat number is associated with expression of nearby genes (eSTRs). Our analysis reveals more than 28,000 eSTRs. We employ fine-mapping to quantify the probability that each eSTR is causal and characterize a group of the top 1,400 fine-mapped eSTRs. We identify hundreds of eSTRs linked with published GWAS signals and implicate specific eSTRs in complex traits including height and schizophrenia, inflammatory bowel disease, and intelligence. Overall, our results support the hypothesis that eSTRs contribute to a range of human phenotypes and will serve as a valuable resource for future studies of complex traits.

## Introduction

Genome-wide association studies (GWAS) have identified thousands of genetic loci associated with complex traits^1^, but determining the causal variants, target genes, and biological mechanisms responsible for each signal has proven challenging. The vast majority of GWAS signals lie in non-coding regions^2^ which are difficult to interpret. Expression quantitative trait loci (eQTL) studies attempt to link regulatory genetic variation to gene expression changes as a potential molecular intermediate that drives biological aberrations leading to disease^3^. Indeed, recent studies have utilized eQTL catalogs to pinpoint causal genes and relevant tissues for a variety of traits^4–6^.

An additional major challenge in interpreting GWAS is that lead variants are rarely causal themselves, but rather tag a set of candidate variants in linkage disequilibrium (LD). A variety of statistical fine-mapping techniques have been developed to identify the most likely causal variant^7–9^ considering factors such as summary association statistics, LD information, and functional annotations. However, these methods have been limited by focusing on bi-allelic single nucleotide polymorphisms (SNPs) or short indels. On the other hand, multiple recent studies to dissect GWAS loci have found complex repetitive^6,10^ and structural variants^11–13^ to be the underlying causal variants, highlighting the need to consider additional variant classes.

Short tandem repeats (STRs), consisting of consecutively repeated units of 1-6bp, represent a large source of genetic variation. STR mutation rates are orders of magnitude higher than SNPs^14^ and short indels^15^ and each individual is estimated to harbor around 100 *de novo* mutations in STRs^16^. Expansions at several dozen STRs have been known for decades to cause Mendelian disorders^17^ including Huntington’s Disease and hereditary ataxias. Importantly, these pathogenic STRs represent a small minority of the more than 1.5 million STRs in the human genome^18^. Due to bioinformatics challenges of analyzing repetitive regions, many STRs are often filtered from genome-wide studies^19^. However, increasing evidence supports a widespread role of common variation at STRs in complex traits such as gene expression^20–23^.

STRs may regulate gene expression through a variety of mechanisms^24^. For example, the CCG repeat implicated in Fragile X Syndrome was shown to disrupt DNA methylation, altering expression of *FMR1*^25^. Yeast studies have demonstrated that homopolymer repeats act as nucleosome positioning signals with downstream regulatory effects^26,27^. Dinucleotide repeats may alter affinity of nearby DNA binding sites^28^. Furthermore, certain STR repeat units may form non-canonical DNA and RNA secondary structures such as G-quadruplexes^29^, R-loops^30^, and Z-DNA^31^.

We previously performed a genome-wide analysis to identify more than 2,000 STRs for which the number of repeats were associated with expression of nearby genes^20^, termed expression STRs (eSTRs). However, the quality of the datasets available for this study reduced power to detect associations and prevented accurate fine-mapping of individual eSTRs. First, STR genotypes were based on low coverage (4-6x) whole genome sequencing data performed using short reads (50-100bp) which are unable to span across many STRs. As a result, individual STR genotype calls exhibited poor quality with less than 50% genotyping accuracy^18^. Second, the study was based on a single cell-type (lymphoblastoid cell lines; LCLs) with potentially limited relevance to most complex traits^32^. While our and other studies^20,22^ demonstrated that eSTRs explain a significant portion (10-15%) of the *cis* heritability of gene expression, the resulting eSTR catalogs were not powered to causally implicate eSTRs over other nearby variants.

Here, we leverage deep whole genome sequencing (WGS) and expression data collected by the Genotype-Tissue Expression Project (GTEx)^33^ to map eSTRs in 17 tissues. Our analysis reveals more than 28,000 unique eSTRs. We employ fine-mapping to quantify the probability of causality of each eSTR and characterize the top 1,400 (top 5%) fine-mapped eSTRs. We additionally identify hundreds of eSTRs that are in strong LD with published GWAS signals and implicate specific eSTRs in multiple complex traits including height, schizophrenia inflammatory bowel disease, and intelligence. To further validate our findings, we employ available GWAS data to demonstrate evidence of a causal link between an eSTR for *RFT1* and height and use a reporter assay to experimentally validate an effect of this STR on expression. Finally, our eSTR catalog is publicly available and can provide a valuable resource for future studies of complex traits.

## Results

### Profiling expression STRs across 17 human tissues

We performed a genome-wide analysis to identify associations between the number of repeats in each STR and expression of nearby genes (expression STRs, or “eSTRs”, which we use to refer to a unique STR by gene association). We focused on 652 individuals from the GTEx^33^ dataset for which both high coverage WGS and RNA-sequencing of multiple tissues were available (Fig. 1a). We used HipSTR^34^ to genotype STRs in each sample. After filtering low quality calls (**Methods**), 175,226 STRs remained for downstream analysis. To identify eSTRs, we performed a linear regression between average STR length and normalized gene expression for each individual at each STR within 100kb of a gene, controlling for sex, population structure, and technical covariates (**Methods**, Supplementary Figs. 1-3). Analysis was restricted to 17 tissues where we had data for at least 100 samples (**Supplementary Table 1, Methods**) and to genes with median RPKM greater than 0. Altogether, we performed an average of 262,593 STR-gene tests across 15,840 protein-coding genes per tissue.

**Figure 1:**
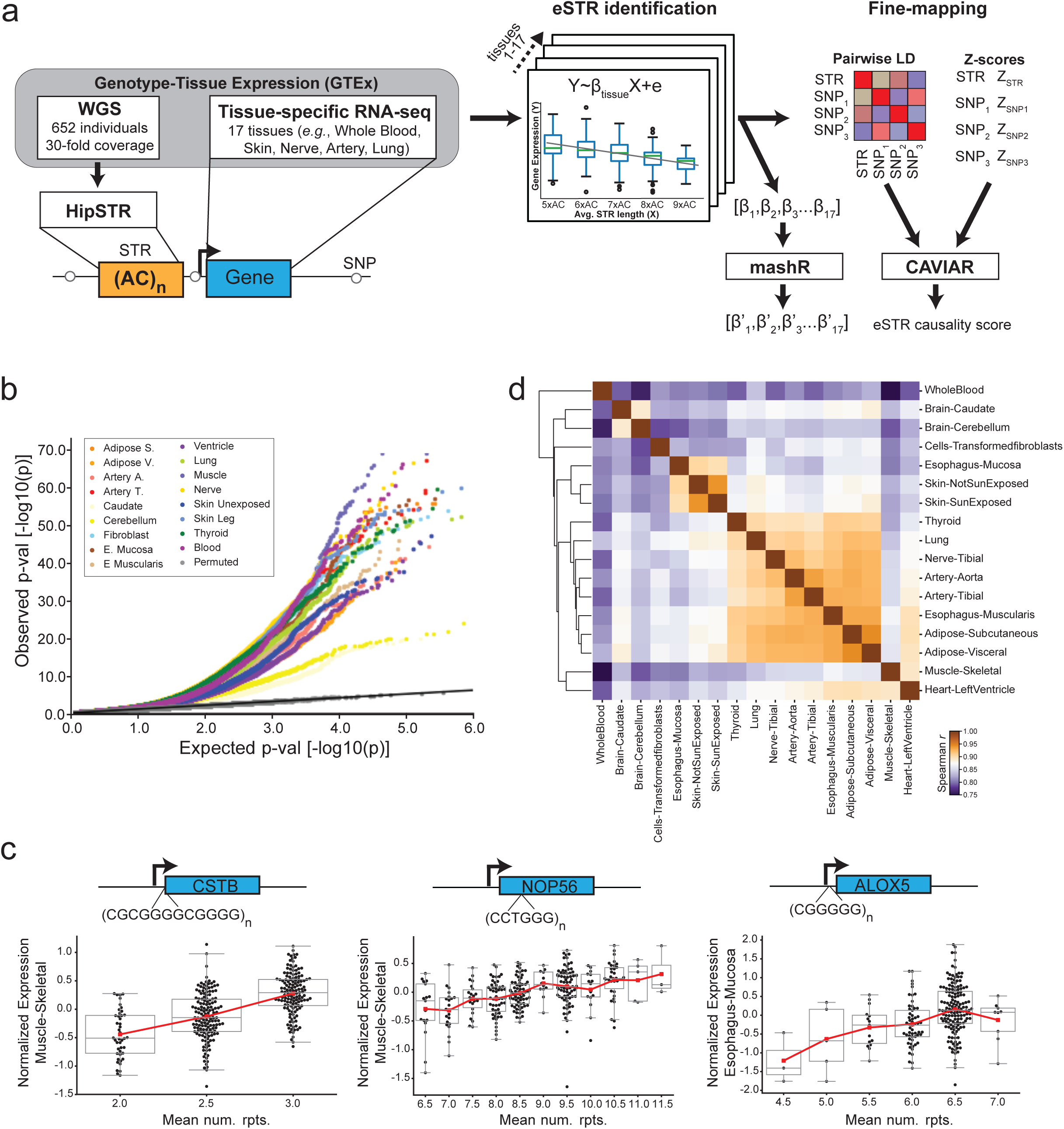
Multi-tissue identification of eSTRs. **(a) Schematic of eSTR discovery pipeline.** We analyzed RNA-seq from 17 tissues and STR genotypes obtained from deep WGS for 652 individuals from the GTEx Project. For each STR within 100kb of a gene, we tested for association between length of the STR and expression of the gene in each tissue. For each gene, CAVIAR was used to fine-map the effects of eSTRs vs. nominally significant *cis* SNPs on gene expression. CAVIAR takes as input pairwise variant LD and effect sizes (Z-scores) and outputs a posterior probability of causality for each variant. For multi-tissue analysis, per-tissue effect sizes and standard errors were used as input to mashR, which computes posterior effect size estimates in each tissue based on potential sharing of eSTRs across tissues. **(b) eSTR association results.** The quantile-quantile plot compares observed p-values for each STR-gene test vs. the expected uniform distribution. Gray dots denote permutation controls. The black line shows the diagonal. Colored dots show observed p-values for each tissue. **(c) Example eSTRs previously implicated in disease.** Left: a CG-rich FM-eSTR upstream of *CSTB* was previously implicated in myoclonus epilepsy^73^. Middle: a multi-allelic intronic CCTGGG FM-eSTR in *NOP56* was implicated in spinocerebellar ataxia 36^74^. Right: A CGGGGG FM-eSTR in the promoter of *ALOX5* was previously shown to regulate *ALOX5* expression in leukocytes^42^ and is associated with reduced lung function^75^ and cardiovascular disease^76^. Each black point represents a single individual. For each plot, the x-axis represents the mean number of repeats in each individual and the y-axis represents normalized expression in a representative tissue. Boxplots summarize the distribution of expression values for each genotype. Horizontal lines show median values, boxes span from the 25th percentile (Q1) to the 75th percentile (Q3). Whiskers extend to Q1-1.5*IQR (bottom) and Q3+1.5*IQR (top), where IQR gives the interquartile range (Q3-Q1). The red line shows the mean expression for each x-axis value. Gene diagrams are not drawn to scale. **(d) eSTR effect sizes are correlated across tissues and studies.** Each cell in the matrix shows the Spearman correlation between mashR FM-eSTR effect sizes (β’) for each pair of tissues. Only eSTRs with CAVIAR score >0.3 in at least one tissue (FM-eSTRs) were included in each correlation analysis. Rows and columns were clustered using hierarchical clustering (**Methods**).

Using this approach, we identified 28,375 unique eSTRs associated with 12,494 genes in at least one tissue at a gene-level FDR of 10% (Fig. 1b, **Supplementary Table 1, Supplementary Dataset 1**). The number of eSTRs detected per tissue correlated with sample size as expected (Pearson r=0.75; p=0.00059; n=17), with the smallest number of eSTRs detected in the two brain tissues presumably due to their low sample sizes (Supplementary Fig. 4). Notably, although whole blood and skeletal muscle had the highest number of samples, we identified fewer eSTRs in those tissues than in others with lower sample sizes. This finding is concordant with previous results for SNPs in this cohort^33^ and may reflect higher cell-type heterogeneity in these tissue samples. eSTR effect sizes previously measured in LCLs were significantly correlated with effect sizes in all GTEx tissues (p<0.01 for all tissues, mean Pearson r=0.45). We additionally examined previously reported eSTRs^35–42^ that were mostly identified using *in vitro* constructs. Six of eight examples were significant eSTRs in GTEx (p<0.01) in at least one tissue analyzed (**Supplementary Table 2**).

eSTRs identified above could potentially be explained by tagging nearby causal variants such as single nucleotide polymorphisms (SNPs). To identify potentially causal eSTRs we employed CAVIAR^7^, a statistical fine-mapping framework for identifying causal variants. CAVIAR models the relationship between LD-structure and association scores of local variants to quantify the posterior probability of causality for each variant (which we refer to as the CAVIAR score). We used CAVIAR to fine-map eSTRs against all SNPs nominally associated (p<0.05) with each gene under our model (**Methods**, Fig. 1a). On average across tissues, 12.2% of eSTRs had the highest causality scores of all variants tested.

We ranked eSTRs by the best CAVIAR score across tissues and chose the top 5% (best CAVIAR score>0.3) for downstream analysis. We hereby refer to this group as fine-mapped eSTRs (FM-eSTRs) (**Supplementary Table 1, Supplementary Dataset 2**). Expected gene annotations are more strongly enriched in this subset of eSTRs compared to the entire set (Supplementary Fig. 5), and stricter thresholds reduced power to detect eSTR-enriched features described below. Of FM-eSTRs in each tissue, on average 78% explained more gene expression variation beyond that of the best SNP (ANOVA q<0.1). Furthermore, on average each FM-eSTR had CAVIAR score 0.41 higher (41% higher posterior probability) than the top-scoring eSNP (Supplementary Fig. 6). Multiple STRs with known disease implications were captured by this list (Fig. 1c). In many cases, FM-eSTRs show clear relationships between the number of repeats and gene expression across a wide range of repeat lengths (Supplementary Fig. 7).

We next analyzed sharing of eSTRs (defined by a unique STR-gene pair) across tissues. To minimize power differences across tissues and enable cross-tissue comparisons of eSTR effects, we applied multivariate adaptive shrinkage (mash^43^) (Fig. 1a). Mash takes as input effect sizes and standard errors as computed above and recomputes posterior estimates of each while considering global correlations of effect sizes across tissues. We computed correlations of mash effect sizes for FM-eSTRs across all pairs of tissues (Fig. 1d) and recovered previously observed relationships^43^. For example, tissues with similar origins (*e.g.*, Adipose-Visceral/Adipose-Subcutaneous) are highly concordant, whereas Whole Blood effects are less correlated with other tissues. These results are also supported by replication between single-tissue eSTRs using unadjusted effect sizes (Supplementary Fig. 8). We further examined tissue sharing of FM-eSTRs by counting the number of tissues for which mash computed a posterior Z-score with absolute value >4. Most eSTRs are either shared across all tissues analyzed or are shared by only a small number of tissues (Supplementary Fig. 9), again similar to previously reported eSNP analyses in this cohort^33^.

### FM-eSTRs demonstrate unique genomic characteristics and sequence features

We next sought to characterize properties of STRs that might provide insights into their biological function. We reasoned that characteristics such as genomic localization, sequence features, and direction of effects that distinguish FM-eSTRs from all analyzed STRs would support the hypothesis that a subset of them are acting as causal variants. We first considered whether the localization of FM-eSTRs differed from that of STRs overall (Fig. 2a-b, Supplementary Fig. 10). Overall, the majority of FM-eSTRs occur in intronic or intergenic regions, and only 11 FM-eSTRs fall in coding exons (**Supplementary Table 3)**. However, compared to all STRs, those closest to TSSs and near DNAseI HS sites were more likely to act as FM-eSTRs (Fig. 2c-d). FM-eSTRs are strongly enriched at 5’ UTRs (OR=5.0; Fisher’s two-sided p=4.9e-13), 3’ UTRs (OR=2.78; p=5.85e-10), and within 3kb of transcription start sites (OR=3.39; p=1.10e-46). These enrichments are considerably stronger for FM-eSTRs compared to all eSTRs (**Supplementary Table 4**), suggesting as expected that FM-eSTRs are more likely to be causal.

**Figure 2:**
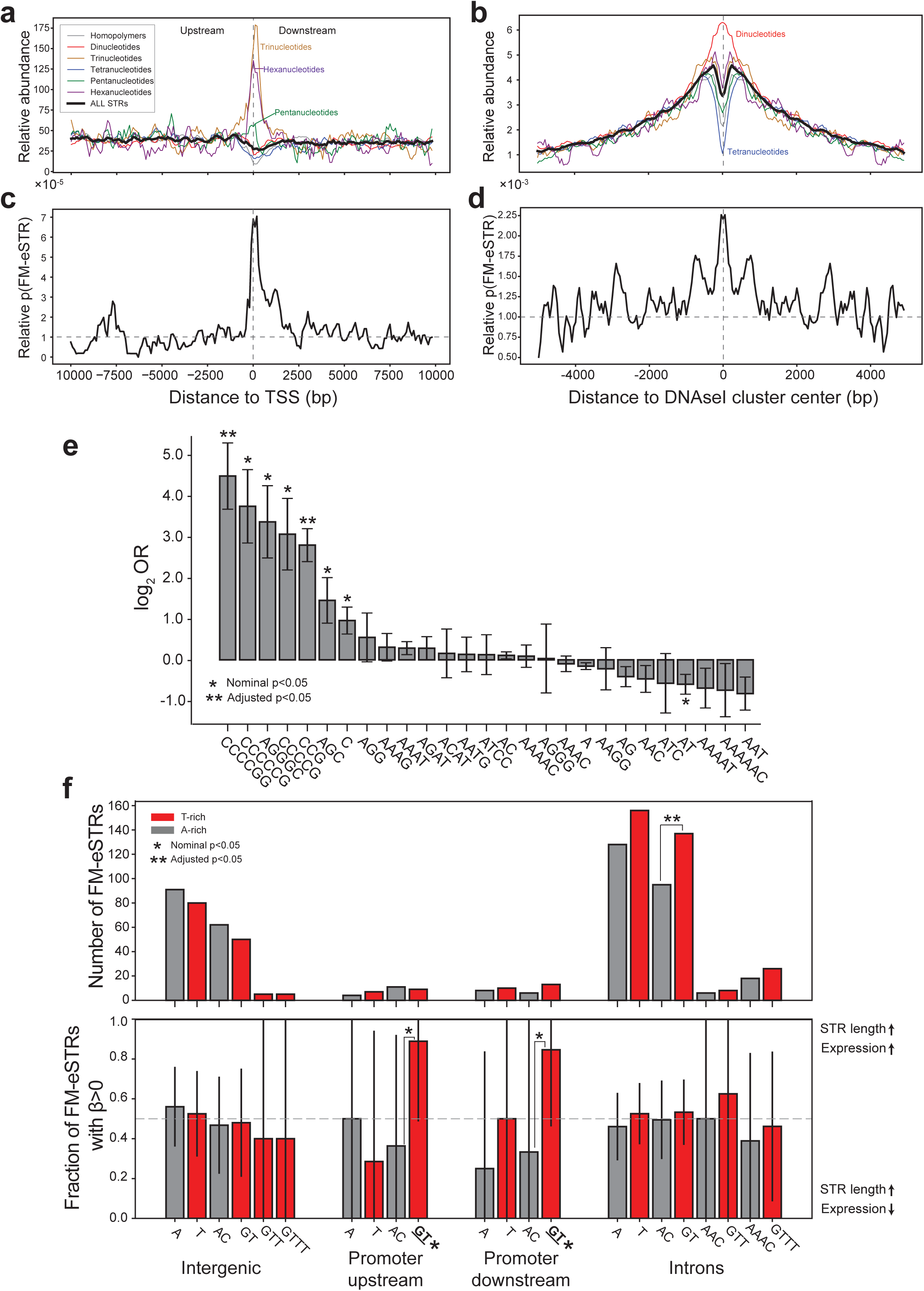
Characterization of FM-eSTRs. **(a) Density of all STRs around transcription start sites.** The y-axis shows the number of STRs in each 100bp bin around the TSS relative to the average across all bins. Negative x-axis numbers denote upstream regions and positive numbers denote downstream regions. **(b) Density of all STRs around DNAseI HS sites.** Plots are centered at ENCODE DNAseI HS clusters and represent the relative number of STRs in each 50bp bin. For (**a)** and (**b)** the black line denotes all STRs and colored lines denote repeats with different repeat unit lengths (gray=homopolymers, red=dinucleotides, gold=trinucleotides, blue=tetranucleotides, green=pentanucleotides, purple=hexanucleotides). **(c) Relative probability to be an FM-eSTR around TSSs**. The black lines represent the probability of an STR in each bin to be an FM-eSTR. Values were scaled relative to the genome-wide average. **(d) Relative probability to be an FM-eSTR around DNAseI HS clusters.** Axes are similar to those in **(c)** except centered around DNAseI HS clusters. For **a-d,** values were smoothed by taking a sliding average of each four consecutive bins. **(e) Repeat unit enrichment at FM-eSTRs across all tissues**. The x-axis shows all repeat units for which there are at least 3 FM-eSTRs across all tissues. The y-axis denotes the log_2_ odds ratios (OR) from performing a Fisher’s exact test comparing FM-eSTRs to all STRs. Enrichments for all eSTRs are given in **Supplementary Table 5**. Single asterisks denote repeat units nominally enriched or depleted (two-sided Fisher exact test p<0.05). Double asterisks denote repeat units significantly enriched after controlling for the number of repeat units tested (Bonferroni adjusted p<0.05). **(f) Strand-biased characteristics of FM-eSTRs.** Top panel: the y-axis shows the number of FM-eSTRs with each repeat unit on the template strand. Bottom panel: the y-axis shows the percentage of FM-eSTRs with each repeat unit on the template strand that have positive effect sizes. Gray bars denote A-rich repeat units (A/AC/AAC/AAAC) and red bars denote T-rich repeat units (T/GT/GTT/GTTT). Single asterisks denote repeat units nominally enriched or depleted (two-sided binomial p<0.05). Double asterisks denote repeat units significantly enriched after controlling for multiple hypothesis testing (Bonferroni adjusted p<0.05). Asterisks above brackets show significant differences between repeat unit pairs. Asterisks on x-axis labels denote departure from the 50% positive effect sizes expected by chance. Error bars give 95% confidence intervals.

To further explore characteristic of FM-eSTRs in regulatory regions, we examined nucleosome occupancy in the lymphoblastoid cell line GM12878 and DNA accessibility measured by DNAseI-seq in a variety of cell types within 500bp of FM-eSTRs (Supplementary Fig. 11). As expected from previous studies^44^, regions near homopolymer repeats are strongly nucleosome-depleted. Notably, STRs with other repeat lengths showed distinct patterns of nucleosome positioning (Supplementary Fig. 11a-c). Nucleosome occupancy is broadly similar for FM-eSTRs compared to all STRs. FM-eSTRs are generally located in regions with higher DNAseI-seq read count compared to non-eSTRs (Mann-Whitney [MW] two-sided p=3.9e-37 in GM12878; Supplementary Fig. 11d-e). DNAseI hypersensitivity around homopolymer FM-eSTRs shows a periodic pattern in GM12878 and other tissue types, with notable peaks located at multiples of 147bp upstream of downstream from the STR (Supplementary Fig. 11d). Given that 147bp is the length of DNA typically wrapped around a single nucleosome^44^, we hypothesize that a subset of homopolymer FM-eSTRs may act by shifting nucleosome positions and thus modulating openness of adjacent hypersensitive sites.

We next examined the sequence characteristics of FM-eSTRs compared to all STRs. We tested FM-eSTRs combined across all tissues for enrichment of each canonical STR repeat unit (defined lexicographically, see **Methods**). FM-eSTRs are most strongly enriched for repeats with GC-rich repeat units (Fig. 2e, **Supplementary Table 5**). For example, the canonical repeat units CCCCGG, CCCCCG, and CCG are 22, 13, and 7-fold enriched in FM-eSTRs compared to all STRs respectively. Notably, the total lengths of FM-eSTRs are significantly higher compared to all STRs analyzed (MW two-sided p=0.00032 and p=2.4e-10 when comparing total repeat number and total length in bp in hg19, respectively).

We next examined effect sizes biases in FM-eSTR associations. Overall, FM-eSTRs are equally likely to show positive vs. negative correlations between repeat length and expression (Supplementary Fig. 12; two-sided binomial p=0.94). We additionally observed that FM-eSTRs with repeat units of the form (A_n_C/G_n_T) show strand-specific effects when in or near transcribed regions. Transcribed FM-eSTRs are more likely to have the T-rich version of the repeat unit on the template strand (two-sided binomial p=0.0015). Further, compared to A-rich repeat units on the template strand, T-rich FM-eSTRs tend to have more positive effect sizes, with the most notable differences for AC vs. GT repeats. These patterns are observed in transcribed regions across multiple distinct repeat types (A/T, AC/GT, AAC/GTT, AAAC/GGGT) but are not present in intergenic regions (Fig. 2f).

Finally, we wondered whether eSTRs might exhibit distinct characteristics in different tissues. We clustered tissue-specific Z-scores (absolute value) for each FM-eSTR calculated jointly across tissues by mash (**Methods**) to identify eight categories of FM-eSTR (Supplementary Fig. 13, 14). These include two clusters of FM-eSTRs present across many tissues (Clusters 2 and 8) as well as several more tissue-specific clusters (*e.g.,* Thyroid for Cluster 1). Notably, clusters do not necessarily imply tissue specificity, but rather enrich for FM-eSTRs with particularly strong effects in one or more tissues compared to others (Supplementary Fig. 14). More than 50% of all genes in each cluster are expressed in all 17 tissues analyzed, and 88% of FM-eSTRs are shared by more than one tissue (Supplementary Fig. 9). Clusters show similar repeat unit enrichment to all FM-eSTRs and do not exhibit distinct enriched repeat units (Supplementary Fig. 15). We further tested whether repeat units of FM-eSTRs are distributed uniformly across clusters. Only one repeat unit (AAAAT) shows a suggestive non-uniform distribution across clusters (Chi-squared p=0.018) with highest prevalence in the thyroid cluster. Similar results were achieved using different numbers of clusters. Overall, our results suggest the majority of eSTRs act by global mechanisms and do not implicate tissue-specific characteristics of FM-eSTRs. However, low numbers of tissue-specific effects limit power to detect differences. Future work may provide insights into potential tissue-specific mechanisms.

### GC-rich eSTRs are predicted to modulate DNA and RNA secondary structure

FM-eSTRs are most strongly enriched for repeats with high GC content (*e.g.*, canonical repeat units CCG, CCCCG, CCCCCG, AGGGC) (Fig. 2e, **Supplementary Table 5**) which are found almost exclusively in promoter regions (Supplementary Fig. 10). These GC-rich repeat units have been shown to form highly stable secondary structures during transcription such as G4 quadruplexes in single-stranded DNA^45^ or RNA^46^ that may regulate gene expression. We hypothesized that the effects of GC-rich eSTRs may be in part due to formation of non-canonical nucleic acid secondary structures that modulate DNA or RNA stability as a function of repeat number. We considered properties of two classes of GC-rich FM-eSTRs: (***i***) those following the standard G4 motif (G_3_N_1-7_G_3_N_1-7_G_3_N_1-7_G_3_)^47^ and (***ii***) repeats with canonical repeat unit CCG which does not meet the standard G4 definition. Notably, the majority of CCG FM-eSTRs (79%) occur in 5’ UTRs compared to only 11% for G4 repeats. We observed that both classes of GC-rich repeats are associated with higher RNAPII (Fig. 3a) and lower nucleosome occupancy (Fig. 3b) compared to all STRs. The relationship with RNAPII was observed across a diverse range of cell and tissue types (Supplementary Fig. 16).

**Figure 3:**
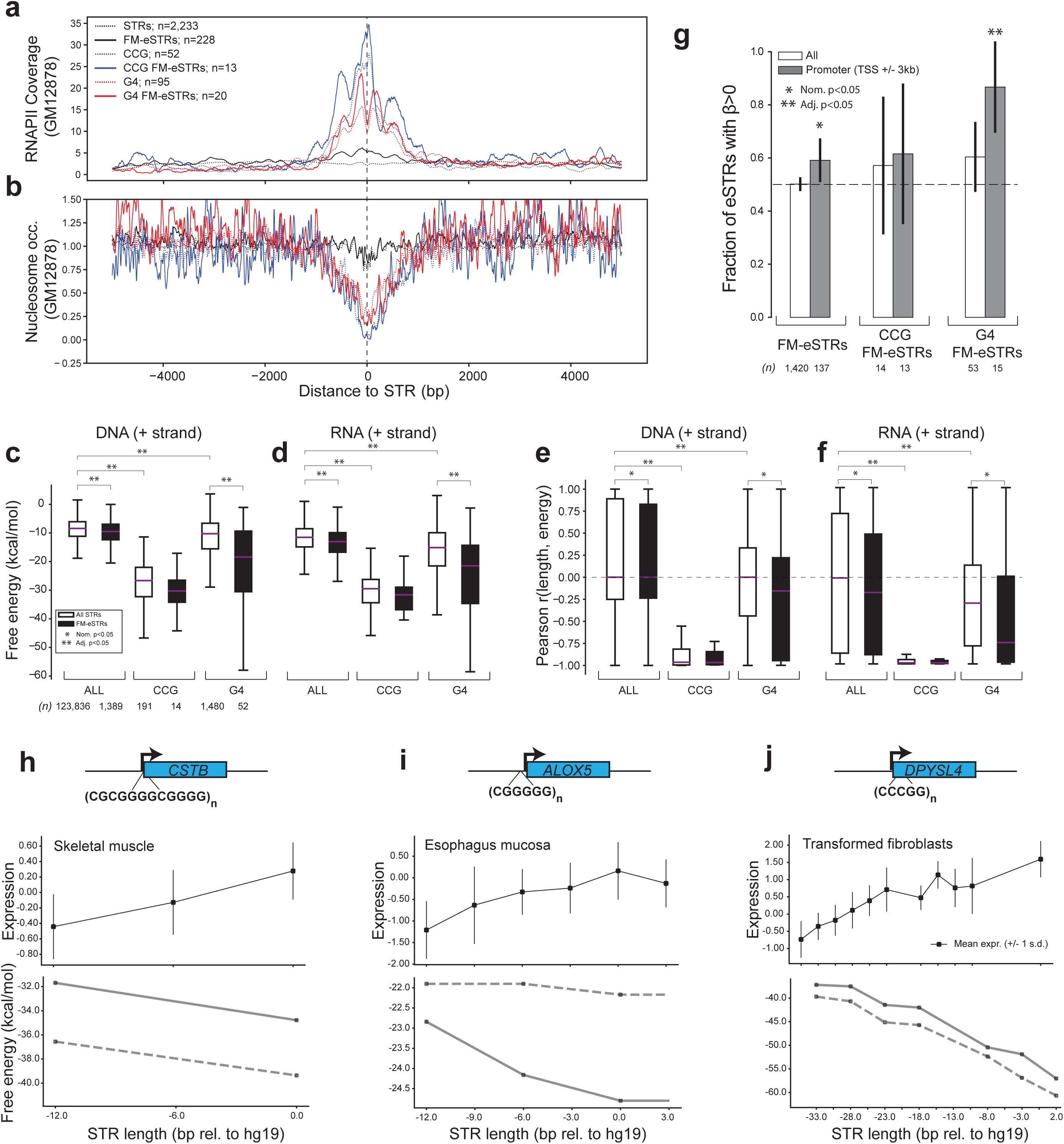
GC-rich eSTRs are predicted to modulate DNA secondary structure. **(a) Density of RNAPII localization around STRs.** The y-axis denotes the average number of ChIP-seq reads for RNA Polymerase II in GM12878 in 5bp bins centered at STRs. **(b) Nucleosome occupancy around STRs.** The y-axis denotes the average nucleosome occupancy in 5bp bins centered at STRs in GM12878. For (**a)** and **(b)**, black lines denote all STRs found within 5kb of TSSs, blue lines denote CCG STRs, and red lines denote STRs matching the canonical G4 motif. Dashed lines represent all STRs of each class and solid lines represent FM-eSTRs. Only STRs within 5,000bp of a TSS are included. **(c-d) Free energy of STR regions.** Boxplots denote the distribution of free energy for each STR +/- 50bp of context sequence, computed as the average across all alleles at each STR. **(c)** and **(d)** show results computed using the template strand for DNA and RNA, respectively. **(e-f) Pearson correlation between STR length and free energy.** Correlations were computed separately for each STR, and plots show the distribution of correlation coefficients across all STRs. The dashed horizontal line denotes 0 correlation as expected by chance. **(e)** and **(f)** show results computed using the template strand for DNA and RNA, respectively. For **c-f**, horizontal purple lines show medians and boxes span from the 25th percentile (Q1) to the 75th percentile (Q3). Whiskers extend to Q1-1.5*IQR (bottom) and Q3+1.5*IQR (top), where IQR represents the interquartile range (Q3-Q1). White boxes show all STRs in each category and black boxes show FM-eSTRs. Upper brackets denote significant differences for all STRs (white) across categories or for significant differences within each category between all STRs (white) and FM-eSTRs (black). Numbers below **(c)** denote the number of data points (unique STRs) included in each box. Nominally significant differences (Mann Whitney one-sided p<0.05) between distributions are denoted with a single asterisk. Differences significant after controlling for multiple hypothesis correction are denoted with double asterisks. For each category (free energy and Pearson correlation), we used a Bonferroni correction to control for 20 total comparisons: comparing all vs. FM-eSTRs separately in each category, comparing CCG vs. all STRs, and comparing G4 vs. all STRs, in four conditions (DNA +/- and RNA +/-). Results for non-template strands are shown in Supplementary Fig. 17. **(g) Bias in the direction of eSTR effect sizes.** The y-axis shows the percentage of FM-eSTRs in each category with positive effect sizes, meaning a positive correlation between STR length and expression. White bars denote all STRs in each category. Gray bars denote STRs falling within 3kb of TSSs. Error bars give 95% confidence intervals. **(h-j) Examples of G4 FM-eSTRs in promoter regions predicted to modulate secondary structure**. For each example, top plots show the mean expression across all individuals with each mean STR length. Vertical bars represent +/- 1 s.d. Bottom plots show the free energy computed by mfold for each number of repeats for the STR. Note, expression plots (top) have additional points to represent heterozygous genotypes, whereas energies (bottom) were computed per-allele rather than per-genotype. Solid and dashed gray lines show energies for alleles on the template and non-template strands, respectively. The x-axis shows STR lengths relative to the hg19 reference genome in bp. Gene diagrams are not drawn to scale.

To evaluate whether GC-rich repeats could be modulating DNA or RNA secondary structure, we used mfold^48^ to calculate the free energy of each STR and 50bp of its surrounding context in single stranded DNA or RNA. We considered all common allele lengths (number of repeats) observed at each STR (**Methods**) and computed energies for both the template and non-template strands. We then computed the correlation between the number of repeats and free energy at each STR region. Overall, both G4 and CGG STRs have lower mean free energy (greater stability) and more negative correlations between repeat number and free energy compared to all STRs (Fig. 3c-f, Supplementary Fig. 17; adjusted MW one-sided p<0.05). Compared to all STRs, FM-eSTRs tend to have lower free energy and more negative correlations with repeat number (MW p<0.05 in all categories except for CGG STRs). Notably, both metrics (mean free energy and correlation of repeat number vs. free energy) are significantly correlated with the total length of STR in all cases (Pearson correlation p<0.01). FM-eSTRs tend to be longer than STRs overall (see above), which may partially explain the secondary structure trends observed.

Based on previous observations^49^, we predicted that higher repeat numbers at GC-rich eSTRs would result in greater DNA or RNA stability and in turn would increase expression of nearby genes. To that end, we tested whether FM-eSTRs were biased toward negative vs. positive effect sizes. As described above, overall FM-eSTRs show no bias in effect direction. However, when considering only repeats in promoter regions (TSS +/- 3kb), 59% of FM-eSTRs have positive effect sizes, significantly more than the 50% expected by chance (binomial two-sided p=0.04; n=137). This effect was stronger when considering only G4 FM-eSTRs (87% positive effect sizes; p=0.0074; n=15) but not significant for CCG FM-eSTRs (62% positive; n=13; p=0.58; Fig. 3g). For multiple G4 FM-eSTRs, expression levels across allele lengths follow an inverse relationship with free energy (Fig. 3h-j). Altogether, these results support a model in which higher repeat numbers at GC-rich eSTRs in promoter regions stabilize DNA secondary structures which promote transcription. Lastly, the contradictory results for CCG STRs may indicate that those repeats could act by distinct mechanisms compared to G4 STRs, but also may be due in part to limited power from a smaller sample size.

### eSTRs are potential drivers of published GWAS signals

Finally, we wondered whether our eSTR catalog could identify STRs affecting complex traits in humans. We first leveraged the NHGRI/EBI GWAS catalog^50^ to identify FM-eSTRs that are nearby and in LD with published GWAS signals. Overall, 1,381 unique FM-eSTRs are within 1Mb of GWAS hits (**Methods, Supplementary Dataset 3**). Of these, 847 are in moderate LD (r^2^>0.1) and 65 are in strong LD (r^2^>0.8) with the lead SNP. For 7 loci in at least moderate LD, the lead GWAS variant is within the STR itself (**Supplementary Table 6**).

We next sought to determine whether specific published GWAS signals could be driven by changes in expression due to an underlying but previously unobserved FM-eSTR. We reasoned that such loci would exhibit the following properties: (***i***) strong similarity in association statistics across variants for both the GWAS trait and expression of a particular gene, indicating the signals may be co-localized, *i.e.*, driven by the same causal variant; and (***ii***) strong evidence that the FM-eSTR causes variation in expression of that gene (Fig. 4a). Co-localization analysis requires high-resolution summary statistic data. Thus, we focused on several example complex traits (height^51^, schizophrenia^52^, inflammatory bowel disease (IBD)^53^, and intelligence^54^) for which detailed summary statistics computed on cohorts of tens of thousands or more individuals are publicly available (**Methods**).

**Figure 4:**
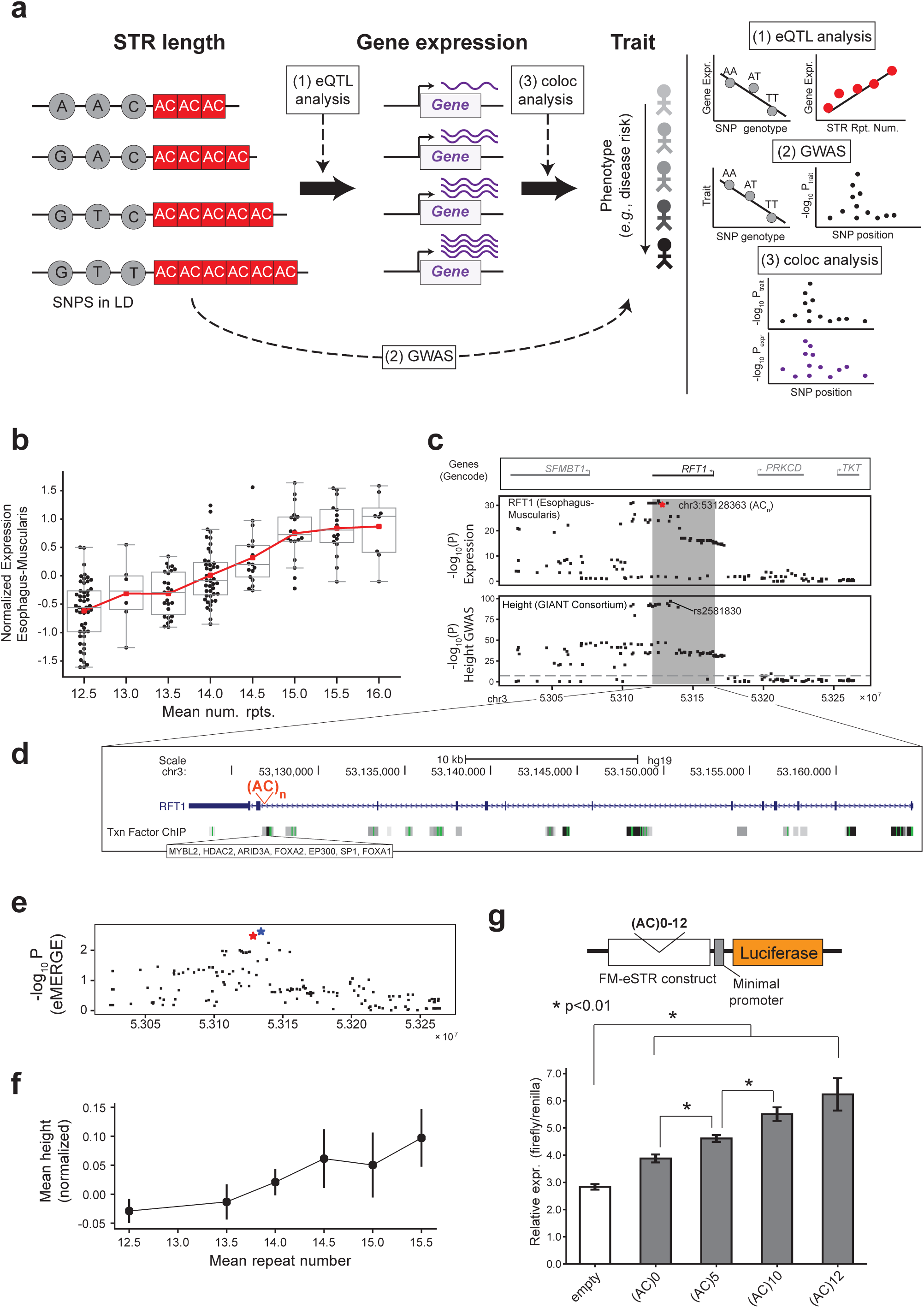
FM-eSTRs co-localize with published GWAS signals. **(a) Overview of analyses to identify FM-eSTRs involved in complex traits.** In cases where FM-eSTRs may drive complex phenotypes, we assumed a model where variation in STR repeat number (red; left) alters gene expression (purple; middle), which in turn affects the value of a particular complex trait (right). Even when the STR is the causal variant, nearby SNPs (gray; left) in LD may be associated with both gene expression through eQTL analysis and the trait of interest through GWAS. Black arrows indicate assumed causal relationships. Dashed arrows indicate analysis approaches (1-3) used to test each relationship. (1) eQTL analysis tests for associations between SNP genotype or STR repeat number and gene expression. (2) GWAS tests for associations between SNP genotype and trait value. (3) coloc analysis tests whether the association signals for expression and the trait are driven by the same underlying causal variant. **(b) eSTR association for *RFT1*.** The x-axis shows STR genotype at an AC repeat (chr3:53128363) as the mean number of repeats and the y-axis gives normalized *RFT1* expression in a representative tissue (Esophagus-Muscularis). Each point represents a single individual. Boxplots summarize expression distributions for each genotype as described in Fig. 1d. Red lines show the mean expression for each x-axis value. **(c) Summary statistics for *RFT1* expression and height.** The top panel shows genes in the region around *RFT1*. The middle panel shows the -log_10_ p-values of association between each variant and *RFT1* expression in aortic artery. The FM-eSTR is denoted by a red star. The bottom panel shows the -log_10_ p-values of association for each variant with height based on available summary statistics^51^. The dashed gray horizontal line shows genome-wide significance threshold. **(d) Detailed view of the *RFT1* locus.** A UCSC genome browser^60^ screenshot is shown for the region in the gray box in **(b)**. The FM-eSTR is shown in red. The bottom track shows transcription factor (TF) binding clusters profiled by ENCODE. **(e) eSTR and SNP associations with height at the *RFT1* locus in the eMERGE cohort.** The x-axis shows the same genomic region as in **(b).** The y-axis denotes association p-values for each variant in the subset of eMERGE cohort samples analyzed here. Black dots represent SNPs. The blue star denotes the top variant in the region identified by Yengo, *et al.*^51^ (rs2581830), which was also the top SNP in eMERGE. The red star represents the imputed FM-eSTR. **(f) Imputed *RFT1* repeat number is positively correlated with height.** The x-axis shows the mean number of AC repeats. The y-axis shows the mean normalized height for all samples included in the analysis with a given genotype. Vertical black lines show +/- 1 s.d. **(g) Reporter assay shows expected positive trend between FM-eSTR repeat number and luciferase expression.** Top: schematic of the experimental design (not to scale). A variable number of AC repeats plus surrounding genomic context (hg19 chr3:53128201-53128577) were introduced upstream of a minimal promoter driving reporter expression. Bottom: white bar shows results from the unmodified plasmid (empty). Gray bars show expression results for constructs with each number of repeats (0, 5, 10, and 12). Reporter expression results normalized to a Renilla control. Error bars show +/- 1 s.d. Asterisks represent significant differences between conditions (one-sided t-test p<0.01).

For each trait, we identified FM-eSTRs within 1Mb of published GWAS signals from **Supplementary Dataset 3.** We then used coloc^55^ to compute the probability that the FM-eSTR signals we derived from GTEx and the GWAS signals derived from other cohorts are co-localized. The coloc tool compares association statistics at each SNP in a region for expression and the trait of interest and returns a posterior probability that the signals are co-localized. We used coloc to test a total of 276 gene×trait pairs (138, 45, 29, and 64 for height, intelligence, IBD, and schizophrenia respectively). In total, we identified 28 GWAS loci with (1) an FM-eSTR in at least moderate LD (r^2^>0.1) with a nearby SNP for that trait in the GWAS catalog and (2) co-localization probability between the target gene and the trait >90% (**Supplementary Table 7**, Supplementary Fig. 18-19).

A top example in our analysis was an FM-eSTR for *RFT1*, an enzyme involved in the pathway of N-glycosylation of proteins^56^, that has 97.8% co-localization probability with a GWAS signal for height (Fig. 4b-c). The lead SNP in the NHGRI catalog (rs2336725) is in high LD (r^2^=0.85) with an AC repeat that is a significant eSTR in 15 tissues. This STR falls in a cluster of transcription factor and chromatin regulator binding regions identified by ENCODE near the 3’ end of the gene (Fig. 4d) and exhibits a positive correlation with expression across a range of repeat numbers.

To more directly test for association between this FM-eSTR and height, we used our recently developed STR-SNP reference haplotype panel^57^ to impute STR genotypes into available GWAS data. We focused on the eMERGE cohort (**Methods**) for which imputed genotype array data and height measurements are available. We tested for association between height and SNPs as well as for AC repeat number after excluding samples with low STR imputation quality (**Methods**). Imputed AC repeat number is significantly associated with height in the eMERGE cohort (p=0.00328; beta=0.010; n=6,393), although with a slightly weaker p-value compared to the top SNP (Fig. 4e). Notably, even in the case that the STR is the causal variant, power is likely reduced due to the lower quality of imputed STR genotypes. Encouragingly, AC repeat number shows a strong positive relationship with height across a range of repeat lengths (Fig. 4f), similar to the relationship between repeat number and *RFT1* expression.

To further investigate whether the FM-eSTR for *RFT1* could be a causal driver of gene expression variation, we devised a dual reporter assay in HEK293T cells to test for an effect of the number of repeats on gene expression (0, 5, 10, or 12 repeats plus approximately 170bp of genomic sequence context on either side (**Supplementary Table 8, Methods**). We observed a positive linear relationship between the number of AC repeats and reporter expression as predicted (Fig. 4g) (Pearson r=0.97; p=0.013). Furthermore, all pairs of constructs with consecutive repeat numbers showed significantly different expression (one-sided t-test p<0.01) with the exception of 10 vs. 12 repeats. Overall, these results further support the hypothesis that eSTRs may act as causal drivers of gene expression.

## Discussion

Here we present the most comprehensive resource of eSTRs to date, which reveals more than 28,000 associations between the number of repeats at STRs and expression of nearby genes across 17 tissues. We performed fine-mapping to quantify the probability that each eSTR causally effects gene expression and characterize top fine-mapped eSTRs. eSTRs analyzed here consist of a large spectrum of repeat classes with a variety of repeat unit lengths and sequences, ranging from homopolymers to hexanucleotide repeats. It is probable that each type induces distinct regulatory effects (Fig. 5). While we explored several potential mechanisms, including nucleosome positioning and the formation of non-canonical DNA or RNA secondary structures, our results do not rule out other potential mechanisms for these eSTRs.

**Figure 5:**
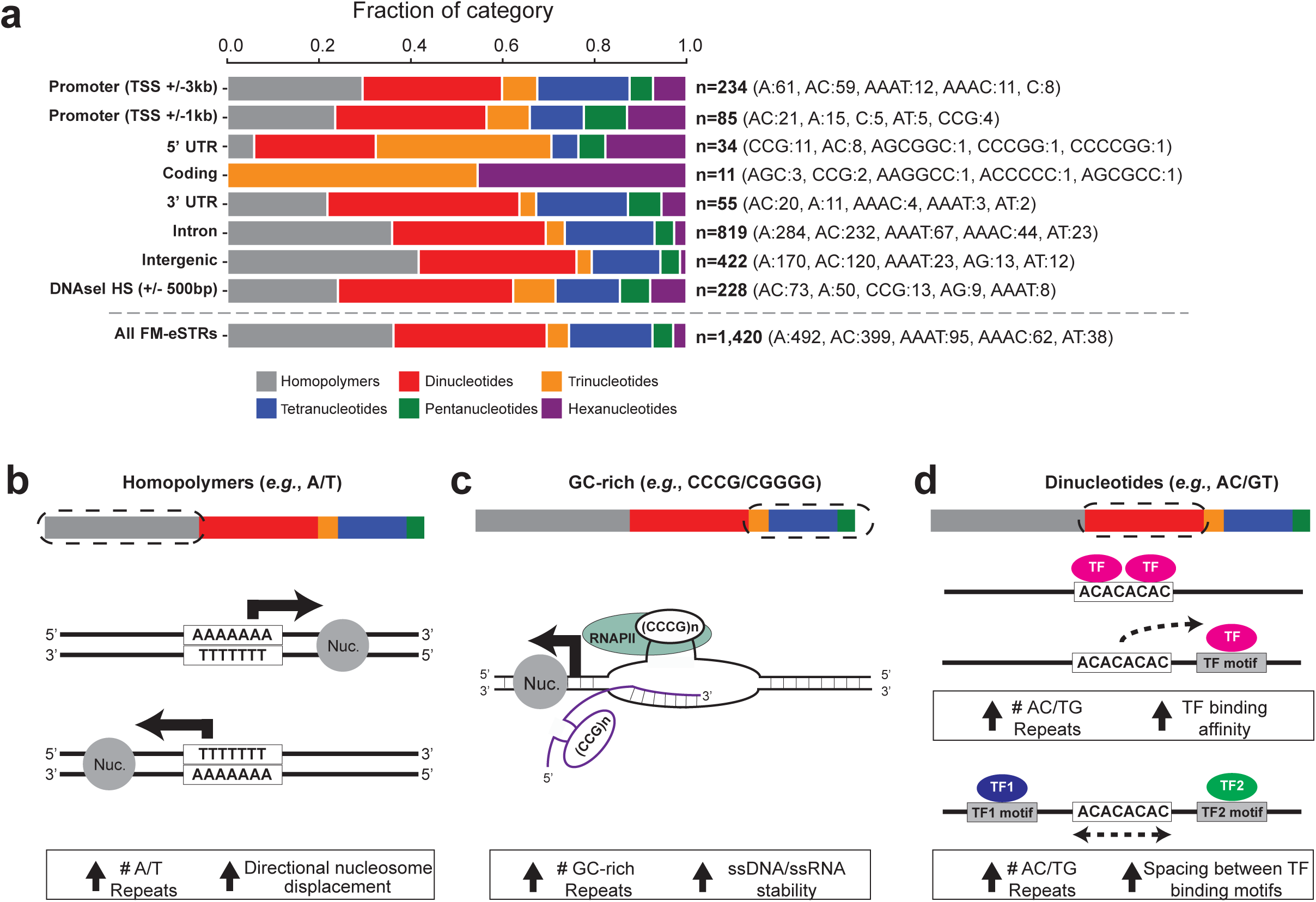
Summary of FM-eSTRs classes and potential regulatory mechanisms. **(a) Distribution of FM-eSTR classes across genomic annotations.** Each bar shows the fraction of FM-eSTRs falling in each annotation consisting of homopolymer (gray), dinucleotide (red), trinucleotide (orange), tetranucleotide (blue), pentanucleotide (green) or hexanucleotide (purple) repeats. The total number of FM-eSTRs and the top five most common repeat units in each category are shown on the right. Note, FM-eSTRs may be counted in more than one category. **(b) Homopolymer A/T STRs are predicted to modulate nucleosome positioning.** Homopolymer repeats are depleted of nucleosomes (gray circles) and may modulate expression changes in nearby genes through altering nucleosome positioning. **(c) GC-rich STRs form DNA and RNA secondary structures during transcription.** Highly stable secondary structures such as G4 quadruplexes may act by expelling nucleosomes (gray circle) or stabilizing RNAPII (light green circle). These structures may form in DNA (black) or RNA (purple) The stability of the structure can depend on the number of repeats. **(d) Dinucleotide STRs can alter transcription factor binding.** Dinucleotides are prevalent in putative enhancer regions. They may potentially alter transcription factor binding by forming binding sites themselves (top), changing affinity of nearby binding sites (middle), or modulating spacing between nearby binding sites (bottom). For **(b)-(d)**, text and arrows in the white boxes provide a summary of the predicted eSTR mechanism depicted in each panel.

We leveraged our resource to provide evidence that FM-eSTRs may drive a subset of published GWAS associations for a variety of complex traits. Altogether, our complex trait analysis demonstrates that STRs may represent an important class of variation that is largely missed by current GWAS. STRs have a unique ability compared to bi-allelic variants such as SNPs or small indels to drive phenotypic variation along a spectrum of multiple alleles, each with different numbers of repeats. In multiple examples, the eSTR shows a linear trend between repeat length and expression across a range of repeat numbers, a signal that cannot be easily explained by tagging nearby bi-allelic variants. Importantly, the cases identified here likely represent a minority of eSTRs driving complex traits. Our analysis is based only on signals that could be detected by standard SNP-based GWAS, which are underpowered to detect underlying multi-allelic associations from STRs^57^. Further work to directly test for associations between STRs and phenotypes is likely to reveal a widespread role for repeat number variation in complex traits.

Our study faced several limitations. (***i***) eSTR discovery was restricted to linear associations between repeat number and expression. Our analysis did not consider non-linear effects or the effect of sequence imperfections, such as SNPs or small indels, within the STR sequence itself. (***ii***) While we applied stringent fine-mapping approaches to find eSTRs whose signals are likely not explained by nearby SNPs in LD, some signals could plausibly be explained by other variant classes such as structural variants^58^ or *Alu* elements^59^ that were not considered. Furthermore, our fine-mapping procedure may be vulnerable to false negatives for STRs in strong or perfect LD with nearby SNPs or false positives due to noise present with small sample sizes. (***iii***) Our study was limited to tissues available from GTEx with sufficient sample sizes. While this greatly expanded on the single tissue used in our previous eSTR analysis, some tissues such as brain were not well represented and had low power for eSTR detection. Further, while we analyzed 17 distinct tissues, due to overwhelming sharing of eSTRs across tissues, we were unable to identify tissue-specific characteristics of eSTRs (***iv***) Despite strong evidence that the FM-eSTRs for *RFT1* and other genes may drive published GWAS signals, we have not definitively proved causality. Additional work is needed to validate effects on expression and evaluate the impact of these STRs in trait-relevant cell types. Nevertheless, most FM-eSTRs have broad effects across many tissue types and in many cases are the most plausible causal variants identified by fine-mapping. This suggests our list of FM-eSTRs colocalized with GWAS signals (**Supplementary Table 7**) will be useful for identifying candidate eSTRs driving complex traits to be explored in future studies.

In summary, our eSTR catalog provides a valuable resource for both obtaining deeper insights into biological roles of eSTRs in regulating gene expression and for identifying potential causal variants underlying a variety of complex traits.

## Materials and Methods

### Dataset and preprocessing

Next-generation sequencing data was obtained from the Genotype-Tissue Expression (GTEx) through dbGaP under phs000424.v7.p2. This included high coverage (30x) Illumina whole genome sequencing (WGS) data and expression data from 652 unrelated individuals (Supplementary Fig. 1). The WGS cohort consisted of 561 individuals with reported European ancestry, 75 of African ancestry, and 8, 3, and 5 of Asian, Amerindian, and Unknown ancestry, respectively. For each sample, we downloaded BAM files containing read alignments to the hg19 reference genome and VCFs containing SNP genotype calls.

STRs were genotyped using HipSTR^34^, which returns the maximum likelihood diploid STR allele sequences for each sample based on aligned reads as input. Samples were genotyped separately with non-default parameters --min-reads 5 and --def-stutter-model. VCFs were filtered using the filter_vcf.py script available from HipSTR using recommended settings for high coverage data (min-call-qual 0.9, max-call-flank-indel 0.15, and max-call-stutter 0.15). VCFs were merged across all samples and further filtered to exclude STRs meeting the following criteria: call rate <80%; STRs overlapping segmental duplications (UCSC Genome Browser^60^ hg19.genomicSuperDups table); penta- and hexamer STRs containing homopolymer runs of at least 5 or 6 nucleotides, respectively in the hg19 reference genome, since we previously found these STRs to have high error rates due to indels in homopolymer regions^57^; and STRs whose frequencies did not meet the percentage of homozygous vs. heterozygous calls based on expected under Hardy-Weinberg Equilibrium (binomial two-sided p<0.05). Additionally, to restrict to polymorphic STRs we filtered STRs with heterozygosity <0.1. Altogether, 175,226 STRs remained for downstream analysis.

We additionally obtained gene-level RPKM values for each tissue from dbGaP project phs000424.v7.p2. We focused on 15 tissues with at least 200 samples, and included two brain tissues with slightly more than 100 samples available (**Supplementary Table 1**). Genes with median RPKM of 0 were excluded and expression values for remaining genes were quantile normalized separately per tissue to a standard normal distribution. Analysis was restricted to protein-coding genes based on GENCODE version 19 (Ensembl 74) annotation.

Prior to downstream analyses, expression values were adjusted separately for each tissue to control for sex, population structure, and technical variation in expression as covariates. For population structure, we used the top 10 principal components resulting from performing principal components analysis (PCA) on the matrix of SNP genotypes from each sample. PCA was performed jointly on GTEx samples and 1000 Genomes Project^61^ samples genotyped using Omni 2.5 genotyping arrays (see **URLs**). Analysis was restricted to bi-allelic SNPs present in the Omni 2.5 data and resulting loci were LD-pruned using plink^62^ with option --indep 50 5 2. PCA on resulting SNP genotypes was performed using smartpca^63,64^. To control for technical variation in expression, we applied PEER factor correction^65^. Based on an analysis of number of PEER factors vs. number of eSTRs identified per tissue (Supplementary Fig. 2), we determined an optimal number of N/10 PEER factors as covariates for each tissue, where N is the sample size. PEER factors were correlated with covariates reported previously for GTEx samples (Supplementary Fig. 3) such as ischemic time.

### eSTR and eSNP identification

For each STR within 100kb of a gene, we performed a linear regression between STR lengths and adjusted expression values:

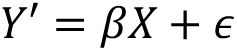

Where *X* denotes STR genotypes, *Y′* denotes expression values adjusted for the covariates described above, *β* denotes the effect size, and ε is the error term. A separate regression analysis was performed for each STR-gene pair in each tissue. For STR genotypes, we used the average repeat length of the two alleles for each individual, where repeat length was computed as a length difference from the hg19 reference, with 0 representing the reference allele. Linear regressions were performed using the OLS function from the Python statsmodels.api module^66^. As a control, for each STR-gene pair we performed a permutation analysis in which sample identifiers were shuffled.

Samples with missing genotypes or expression values were removed from each regression analysis. To reduce the effect of outlier STR genotypes, we removed samples with genotypes observed in less than 3 samples. If after filtering samples there were less than three unique genotypes, the STR was excluded from analysis. Adjusted expression values and STR genotypes for remaining samples were then Z-scaled to have mean 0 and variance 1 before performing each regression. This step forces resulting effect sizes to be between −1 and 1.

We used a gene-level FDR threshold (described previously^20^) of 10% to identify significant STR-gene pairs. We assume most genes have at most a single causal eSTR. For each gene, we determined the STR association with the strongest P-value. This P-value was adjusted using a Bonferroni correction for the number of STRs tested per gene to give a P-value for observing a single eSTR association for each gene. We then used the list of adjusted P-values (one per gene) as input to the fdrcorrection function in the statsmodels.stats.multitest module to obtain a q-value for the best eSTR for each gene. FDR analysis was performed separately for each tissue.

eSNPs were identified using the same model covariates, and normalization procedures but using SNP dosages (0, 1, or 2) rather than STR lengths. Similar to the STR analysis, we removed samples with genotypes occurring in fewer than 3 samples and removed SNPs with less than 3 unique genotypes remaining after filtering. On average, we tested 17 STRs and 533 SNPs per gene.

### Fine-mapping eSTRs

We used model comparison as an orthogonal validation to CAVIAR findings to determine whether the best eSTR for each gene explained variation in gene expression beyond a model consisting of the best eSNP. For each gene with an eSTR we determined the eSNP with the strongest p-value. We then compared two linear models: Y’∼eSNP (SNP-only model) vs. Y’∼eSNP+eSTR (SNP+STR model) using the anova_lm function in the python statsmodels.api.stats module. Q-values were obtained using the fdrcorrection function in the statsmodels.stats.multitest module. On average across tissues, 17.4% of eSTRs tested improved the model over the best eSNP for the target gene (10% FDR). When restricting to FM-eSTRs, 78% improved the model (10% FDR).

We used CAVIAR^7^ v2.2 to further fine-map eSTR signals against the all nominally significant eSNPs (p<0.05) within 100kb of each gene. On average, 121 SNPs per gene passed this threshold and were included in CAVIAR analysis. Pairwise-LD between the eSTR and eSNPs was estimated using the Pearson correlation between SNP dosages (0, 1, or 2) and STR genotypes (average of the two STR allele lengths) across all samples. CAVIAR was run with parameters -f 1 -c 2 to model up to two independent causal variants per locus. In some cases, initial association statistics for SNPs and STRs might have been computed using different sets of samples if some were filtered due to outlier genotypes. To provide a fair comparison between eSTRs and eSNPs, for each CAVIAR analysis we recomputed Z-scores for eSTRs and eSNPs using the same set of samples prior to running CAVIAR.

### Multi-tissue eSTR analysis

We used an R implementation of mash^43^ (mashR) v0.2.21 to compute posterior estimates of eSTR effect sizes and standard errors across tissues. Briefly, mashR takes as input effect sizes and standard error measurements per-tissue, learns various covariance matrices of effect sizes between tissues, and outputs posterior estimates of effect sizes and standard errors accounting for global patterns of effect size sharing. We used all eSTRs with a nominal p-value of <1e-5 in at least one tissue as a set of strong signals to compute covariance matrices. eSTRs that were not analyzed in all tissues were excluded from this step. We included “canonical” covariance matrices (identity matrix and matrices representing condition-specific effects) and matrices learned by extreme deconvolution initialized using PCA with 5 components as suggested by mashR documentation. After learning covariance matrices, we applied mashR to estimate posterior effect sizes and standard errors for each eSTR in each tissue. For eSTRs that were filtered from one or more tissues in the initial regression analysis, we set input effect sizes to 0 and standard errors to 10 in those tissues to reflect high uncertainty in effect size estimates at those eSTRs. For Fig. 1d, rows and columns of the effect size correlation matrix were clustered using default parameters from the clustermap function in the Python seaborn library (see **URLs**).

### Canonical repeat units

For each STR, we defined the canonical repeat unit as the lexicographically first repeat unit when considering all rotations and strand orientations of the repeat sequence. For example, the canonical repeat unit for the repeat sequence CAGCAGCAGCAG would be AGC.

### Enrichment analyses

Enrichment analyses were performed using a two-sided Fisher’s exact test as implemented in the fisher_exact function of the python package scipy.stats (see **URLs**). Overlapping STRs with each annotation was performed using the intersectBed tool of the BEDTools^67^ suite. Genomic annotations were obtained by downloading custom tables using the UCSC Genome Browser^60^ table browser tool to select either coding regions, introns, 5’UTRs, 3’UTRs, or regions upstream of TSSs. An STR could be assigned to more than one category in the case of overlapping transcripts. STRs not assigned to one of those categories were labeled as intergenic. ENCODE DNAseI HS clusters were downloaded from the UCSC Genome Browser (see **URLs**). Analysis was restricted to DNAseI HS clusters annotated in at least 20 cell types. The distance between each STR and the center of the nearest DNAseI HS cluster was computed using the closestBed tool from the BEDTools suite.

### Analysis of DNAseI-seq, ChIP-seq, and Nucleosome occupancy

Genome-wide nucleosome occupancy signal in GM12878 was downloaded from the UCSC Genome Browser (see **URLs**). ChIP-seq reads for RNAPII and DNAseI-seq reads were downloaded from the ENCODE Project website (see **URLs**) (Accessions GM12878 RNAPII: ENCFF775ZJX, heart RNAPII: ENCFF643EGO, lung RNAPII: ENCSR033NHF, tibial nerve RNAPII: ENCFF750HDH, human embryonic stem cells RNAPII: ENCFF526YGE; GM12878 DNAseI: ENCFF775ZJX, fat DNAseI: ENCFF880CAD, tibial nerve DNAseI: ENCFF226ZCG, skin DNAseI: ENCFF238BRB). Histograms of aggregate read densities and heatmaps for individual STR regions were generated using the annoatePeaks.pl tool of Homer^68^. For nucleosome occupancy and DNAseI analyses on all STRs, we used parameters -size 1000 -hist 1. For analysis of GC-rich repeats in promoters, we used parameters -size 10000 -hist 5.

### Characterization of tissue-specific eSTRs

We clustered FM-eSTRs based on effect sizes Z-scores computed by mash for each eSTR in each tissue. We first created a tissue by FM-eSTR matrix of the absolute value of the Z-scores. We then Z-normalized Z-scores for each FM-eSTR to have mean 0 and variance 1. We used the KMeans class from the Python sklearn.cluster module to perform K-means clustering with K=8. The number of clusters was chosen by visualizing the sum of squared distances from centroids for values of K ranging from 1 to 20 and choosing a value of K based on the “elbow method”. Using different values of K produced similar groups. We tested for non-uniform distributions of FM-eSTR repeat units across clusters using a chi-squared test implemented in the scipy.stats chi2_contingency function.

### Analysis of DNA and RNA secondary structure

For each STR, we extracted the repeat plus 50bp flanking sequencing from the hg19 reference genome. We additionally created sequences containing each common allele for each STR. Common alleles were defined as those seen at least 5 times in a previously generated deep catalog of STR variation in 1,916 samples^57^. For each sequence and its reverse complement, we ran mfold^48^ on the DNA and corresponding RNA sequences with mfold arguments NA=DNA and NA=RNA, respectively, and otherwise default parameters to estimate the free energy of each single-stranded sequence. Mann-Whitney tests were performed using the mannwhitneyu function of the scipy.stats python package (see **URLs**).

### Co-localization of FM-eSTRs with published GWAS signals

Published GWAS associations were obtained from the NHGRI/EBI GWAS catalog available from the UCSC Genome Browser Table Browser (table hg19.gwasCatalog) downloaded on July 24, 2019. Height GWAS summary statistics were downloaded from the GIANT Consortium website (see **URLs**). Schizophrenia GWAS summary statistics were downloaded from the Psychiatric Genomics Consortium website (see **URLs**). IBD summary statistics were downloaded from the International Inflammatory Bowel Disease Genetics Consortium (IIBDGC) website. We used the file EUR.IBD.gwas_info03_filtered.assoc with summary statistics in Europeans (see **URLs**). Intelligence summary statistics were downloaded from the Complex Trait Genomics lab website (see **URLs**). LD between STRs and SNPs was computed by taking the squared Pearson correlation between STR lengths and SNP dosages in GTEx samples for each STR-SNP pair. STR genotypes seen less than 3 times were filtered from LD calculations.

Co-localization analysis of eQTL and GWAS signals was performed using the coloc.abf function of the coloc^55^ package. For all traits, dataset 1 was specified as type=”quant” and consisted of eSNP effect sizes and their variances as input. We specified sdY=1 since expression was quantile normalized to a standard normal distribution. Dataset 2 was specified differently for height and schizophrenia to reflect quantitative vs. case-control analyses. For height and intelligence, we specified type=”quant” and used effect sizes and their variances as input. We additionally specified minor allele frequencies listed in the published summary statistics file and the total sample size of N=695,647 and N=269,720 for height and intelligence, respectively. For schizophrenia and IBD, we specified type=”CC” and used effect sizes and their variances as input. We additionally specified the fraction of cases as 33%.

Capture Hi-C interactions (Supplementary Fig. 19) were visualized using the 3D Genome Browser^69^. The visualization depicts interactions profiled in GM12878^70^ and only shows interactions overlapping the STR of interest.

### Association analysis in the eMERGE cohort

We obtained SNP genotype array data and imputed genotypes from dbGaP accessions phs000360.v3.p1 and phs000888.v1.p1 from consent groups c1 (Health/Medical/Biomedical), c3 (Health/Medical/Biomedical - Genetic Studies Only - No Insurance Companies), and c4 (Health/Medical/Biomedical - Genetic Studies Only). Height data was available for samples in cohorts c1 (phs000888.v1.pht004680.v1.p1.c1), c3 (phs000888.v1.pht004680.v1.p1.c3), and c4 (phs000888.v1.pht004680.v1.p1.c4). We removed samples without age information listed. If height was collected at multiple times for the same sample, we used the first data point listed.

Genotype data was available for 7,190, 6100, and 3,755 samples from the c1, c3, and c4 cohorts respectively (dbGaP study phs000360.v3.p1). We performed PCA on the genotypes to infer ancestry of each individual. We used plink to restrict to SNPs with minor allele frequency at least 10% and with genotype frequencies expected under Hardy-Weinberg Equilibrium (p>1e-4). We performed LD pruning using the plink option --indep 50 5 1.5 and used pruned SNPs as input to PCA analysis. We visualized the top two PCs and identified a cluster of 14,147 individuals overlapping samples with annotated European ancestry. We performed a separate PCA using only the identified European samples and used the top 10 PCs as covariates in association tests.

A total of 11,587 individuals with inferred European ancestry had both imputed SNP genotypes and height and age data available. Samples originated from cohorts at Marshfield Clinic, Group Health Cooperative, Northwestern University, Vanderbilt University, and the Mayo Clinic. We adjusted height values by regressing on top 10 ancestry PCs, age, and cohort. Residuals were inverse normalized to a standard normal distribution. Adjustment was performed separately for males and females.

Imputed genotypes (from dbGaP study phs000888.v1.p1) were converted from IMPUTE2^71^ to plink’s binary format using plink, which marks calls with uncertainty >0.1 (score<0.9) as missing. SNP associations were performed using plink with imputed genotypes as input and with the “linear” option with analysis restricted to the region chr3:53022501-53264470.

The *RFT1* FM-eSTR was imputed into the imputed SNP genotypes using Beagle 5^72^ with option gp=true and using our SNP-STR reference haplotype panel^57^. We previously estimated imputation concordance of 97% at this STR in a separate European cohort. Samples with imputed genotype probabilities of less than 0.9 were removed from the STR analysis. We additionally restricted analysis to STR genotypes present in at least 100 samples to minimize the effect of outlier genotypes. We regressed STR genotype (defined above as the average of an individual’s two repeat lengths) on residualized height values for the remaining 6,393 samples using the Python statsmodels.regression.linear_model.OLS function (see **URLs**).

### Dual luciferase reporter assay

Constructs for 0, 5, or 10 copies of AC at the FM-eSTR for *RFT1* (chr3:53128363-53128413 plus approximately 170bp genomic context on either side (RFT1_0rpt, RFT1_5rpt, RFT1_10rpt in **Supplementary Table 8**) were ordered as gBlocks from Integrated DNA technologies (IDT). Each construct additionally contained homology arms for cloning into pGL4.27 (below). We additionally PCR amplified the region from genomic DNA for sample NA12878 with 12 copies of AC (NIGMS Human Genetic Repository, Coriell) using PrimeSTAR max DNA Polymerase (Clontech R045B) and primers RFT1eSTR_F and RFT1eSTR_R (**Supplementary Table 8**) which included the same homology arms.

Constructs were cloned into plasmid pGL4.27 (Promega, E8451), which contains the firefly luciferase coding sequence and a minimal promoter. The plasmid was linearized using EcoRV (New England Biolabs, R3195) and purified from agarose gel (Zymo Research, D4001). Constructs were cloned into the linearized vector using In-Fusion (Clontech, 638910). Sanger sequencing of isolated clones for each plasmid validated expected repeat numbers in each construct.

Plasmids were transfected into the human embryonic kidney 293 cell line (HEK293T; ATCC CRL-3216) and grown in DMEM media (Gibco, 10566-016), supplemented with 10% fetal bovine serum (Gibco, 10438-026), 2 mM glutamine (Gibco, A2916801), 100 units/mL of penicillin, 100 µg/mL of streptomycin, and 0.25 µg/mL Amphotericin B (Anti-Anti Gibco, 15240062). Cells were maintained at 37°C in a 5% CO_2_ incubator. 2×10^5^ HEK293T cells were plated onto each well of a 25 ug/ml poly-D lysine (EMD Millipore, A-003-E) coated 24-well plate, the day prior to transfection. On the day of the transfection medium was changed to Opti-MEM. We conducted co-transfection experiments to test expression of each construct. 100ng of the empty pGL4.27 vector (Promega, E8451) or 100 ng of each one of the pGL4.27 derivatives, were mixed with 5ng of the reference plasmid, pGL4.73 (Promega, E6911), harboring SV40 promoter upstream of Renilla luciferase, and added to the cells in the presence of Lipofectamine™ 3000 (Invitrogen, L3000015), according to the manufacturer’s instructions. Cells were incubated for 24 hr at 37°C, washed once with phosphate-buffered saline, and then incubated in fresh completed medium for an additional 24 hr.

48 hours after transfection the HEK293T cells were washed 3 times with PBS and lysed in 100μl of Passive Lysis Buffer (Promega, E1910). Firefly luciferase and Renilla luciferase activities were measured in 10μl of HEK293T cell lysate using the Dual-Luciferase Reporter assay system (Promega, E1910) in a Veritas™ Microplate Luminometer. Relative activity was defined as the ratio of firefly luciferase activity to Renilla luciferase activity. For each plasmid, transfection and the expression assay were done in triplicates using three wells of cultured cells that were independently transfected (biological repeats), and three individually prepared aliquots of each transfection reaction (technical repeats). Values from each technical replicate were averaged to get one ratio for each biological repeat. Values shown in Fig. 4g represent the mean and standard deviation across the three biological replicates for each construct.

## Data Availability

All eSTR summary statistics are available for download on WebSTR http://webstr.ucsd.edu/downloads.

## Code Availability

Code for performing analyses and generated figures is available at http://github.com/gymreklab/gtex-estrs-paper.

## URLs

1000 Genomes phased Omni2.5 SNP data, ftp.1000genomes.ebi.ac.uk/vol1/ftp/release/20130502/supporting/shapeit2_scaffolds/hd_chip_scaffolds/

ENCODE DNAseI HS clusters, http://hgdownload.cse.ucsc.edu/goldenpath/hg19/encodeDCC/wgEncodeRegDnaseClustered/wgEncodeRegDnaseClusteredV3.bed.gz

ChIP-seq and DNAseI-seq, https://www.encodeproject.org

Nucleosome occupancy in GM12878, http://hgdownload.cse.ucsc.edu/goldenpath/hg19/encodeDCC/wgEncodeSydhNsome/wgEncodeSydhNsomeGm12878Sig.bigWig

Height GWAS summary statistics, https://portals.broadinstitute.org/collaboration/giant/images/0/0f/Meta-analysis_Locke_et_al%2BUKBiobank_2018.txt.gz

Schizophrenia GWAS summary statistics, https://www.med.unc.edu/pgc/results-and-downloads

IBD summary statistics, ftp://ftp.sanger.ac.uk/pub/consortia/ibdgenetics/iibdgc-trans-ancestry-filtered-summary-stats.tgz

Intelligence summary statistics, https://ctg.cncr.nl/documents/p1651/SavageJansen_IntMeta_sumstats.zip

Python scipy.stats package, https://docs.scipy.org/doc/scipy/reference/stats.html

Python statsmodels package, https://www.statsmodels.org

Python scikit-learn package, https://scikit-learn.org/stable/

Python seaborn package, https://seaborn.pydata.org/

MashR vignettes, https://stephenslab.github.io/mashr/articles/intro_mash_dd.html#data-driven-covariances

## Acknowledgements

Research reported in this publication was supported in part by the Office Of The Director, National Institutes of Health under Award Number DP5OD024577. We thank Vineet Bafna, Eric Mendenhall, Joseph Gleeson, and Yu-Ting Liu for helpful comments.

## GTEx Project

The Genotype-Tissue Expression (GTEx) Project was supported by the Common Fund of the Office of the Director of the National Institutes of Health (commonfund.nih.gov/GTEx). Additional funds were provided by the NCI, NHGRI, NHLBI, NIDA, NIMH, and NINDS. Donors were enrolled at Biospecimen Source Sites funded by NCI\Leidos Biomedical Research, Inc. subcontracts to the National Disease Research Interchange (10XS170), GTEx Project March 5, 2014 version Page 5 of 8 Roswell Park Cancer Institute (10XS171), and Science Care, Inc. (X10S172). The Laboratory, Data Analysis, and Coordinating Center (LDACC) was funded through a contract (HHSN268201000029C) to the The Broad Institute, Inc. Biorepository operations were funded through a Leidos Biomedical Research, Inc. subcontract to Van Andel Research Institute (10ST1035). Additional data repository and project management were provided by Leidos Biomedical Research, Inc. (HHSN261200800001E). The Brain Bank was supported supplements to University of Miami grant DA006227. Statistical Methods development grants were made to the University of Geneva (MH090941 & MH101814), the University of Chicago (MH090951,MH090937, MH101825, & MH101820), the University of North Carolina - Chapel Hill (MH090936), North Carolina State University (MH101819),Harvard University (MH090948), Stanford University (MH101782), Washington University (MH101810), and to the University of Pennsylvania (MH101822). The datasets used for the analyses described in this manuscript were obtained from dbGaP at http://www.ncbi.nlm.nih.gov/gap through dbGaP accession number phs000424.v7.p2.

## eMERGE Project - Mayo Clinic

Samples and associated genotype and phenotype data used in this study were provided by the Mayo Clinic. Funding support for the Mayo Clinic was provided through a cooperative agreement with the National Human Genome Research Institute (NHGRI), Grant #: UOIHG004599; and by grant HL75794 from the National Heart Lung and Blood Institute (NHLBI). Funding support for genotyping, which was performed at The Broad Institute, was provided by the NIH (U01HG004424). The datasets used for analyses described in this manuscript were obtained from dbGaP at http://www.ncbi.nlm.nih.gov/gap through dbGaP accession numbers phs000360.v3.p1 and phs000888.v1.p1.

## eMERGE Project - Marshfield Clinic

Funding support for the Personalized Medicine Research Project (PMRP) was provided through a cooperative agreement (U01HG004608) with the National Human Genome Research Institute (NHGRI), with additional funding from the National Institute for General Medical Sciences (NIGMS) The samples used for PMRP analyses were obtained with funding from Marshfield Clinic, Health Resources Service Administration Office of Rural Health Policy grant number D1A RH00025, and Wisconsin Department of Commerce Technology Development Fund contract number TDF FYO10718. Funding support for genotyping, which was performed at Johns Hopkins University, was provided by the NIH (U01HG004438). The datasets used for analyses described in this manuscript were obtained from dbGaP at http://www.ncbi.nlm.nih.gov/gap through dbGaP accession numbers phs000360.v3.p1 and phs000888.v1.p1.

## eMERGE Project - Northwestern University

Samples and data used in this study were provided by the NUgene Project (www.nugene.org). Funding support for the NUgene Project was provided by the Northwestern University’s Center for Genetic Medicine, Northwestern University, and Northwestern Memorial Hospital. Assistance with phenotype harmonization was provided by the eMERGE Coordinating Center (Grant number U01HG04603). This study was funded through the NIH, NHGRI eMERGE Network (U01HG004609). Funding support for genotyping, which was performed at The Broad Institute, was provided by the NIH (U01HG004424). The datasets used for analyses described in this manuscript were obtained from dbGaP at http://www.ncbi.nlm.nih.gov/gap through dbGaP accession numbers phs000360.v3.p1 and phs000888.v1.p1.

## eMERGE Project - Vanderbilt University

Funding support for the Vanderbilt Genome-Electronic Records (VGER) project was provided through a cooperative agreement (U01HG004603) with the National Human Genome Research Institute (NHGRI) with additional funding from the National Institute of General Medical Sciences (NIGMS). The dataset and samples used for the VGER analyses were obtained from Vanderbilt University Medical Center’s BioVU, which is supported by institutional funding and by the Vanderbilt CTSA grant UL1RR024975 from NCRR/NIH. Funding support for genotyping, which was performed at The Broad Institute, was provided by the NIH (U01HG004424). The datasets used for analyses described in this manuscript were obtained from dbGaP at http://www.ncbi.nlm.nih.gov/gap through dbGaP accession numbers phs000360.v3.p1 and phs000888.v1.p1.

## eMERGE Project - Group Health Cooperative

Funding support for Alzheimer’s Disease Patient Registry (ADPR) and Adult Changes in Thought (ACT) study was provided by a U01 from the National Institute on Aging (Eric B. Larson, PI, U01AG006781). A gift from the 3M Corporation was used to expand the ACT cohort. DNA aliquots sufficient for GWAS from ADPR Probable AD cases, who had been enrolled in Genetic Differences in Alzheimer’s Cases and Controls (Walter Kukull, PI, R01 AG007584) and obtained under that grant, were made available to eMERGE without charge. Funding support for genotyping, which was performed at Johns Hopkins University, was provided by the NIH (U01HG004438). Genome-wide association analyses were supported through a Cooperative Agreement from the National Human Genome Research Institute, U01HG004610 (Eric B. Larson, PI). The datasets used for analyses described in this manuscript were obtained from dbGaP at http://www.ncbi.nlm.nih.gov/gap through dbGaP accession numbers phs000360.v3.p1 and phs000888.v1.p1.

## Author Contributions

S.F.F. performed all eSTR and eSNP mapping, helped perform downstream analyses and helped draft the manuscript. J.M. performed multi-tissue analysis using mashR and helped revise the manuscript. C.W. optimized and performed the reporter assay. S.S. participated in design of the STR imputation analysis. S.S.-B. lead, designed, and analyzed data from the reporter assay. R.Y. implemented the WebSTR web application. A.G. conceived and planned analyses and validation experiments of regulatory effects of eSTRs and wrote the manuscript. M.G. conceived the study, designed and performed analyses, and drafted the initial manuscript. All authors have read and approved the final manuscript.

---

**Supplementary Figure 1:**
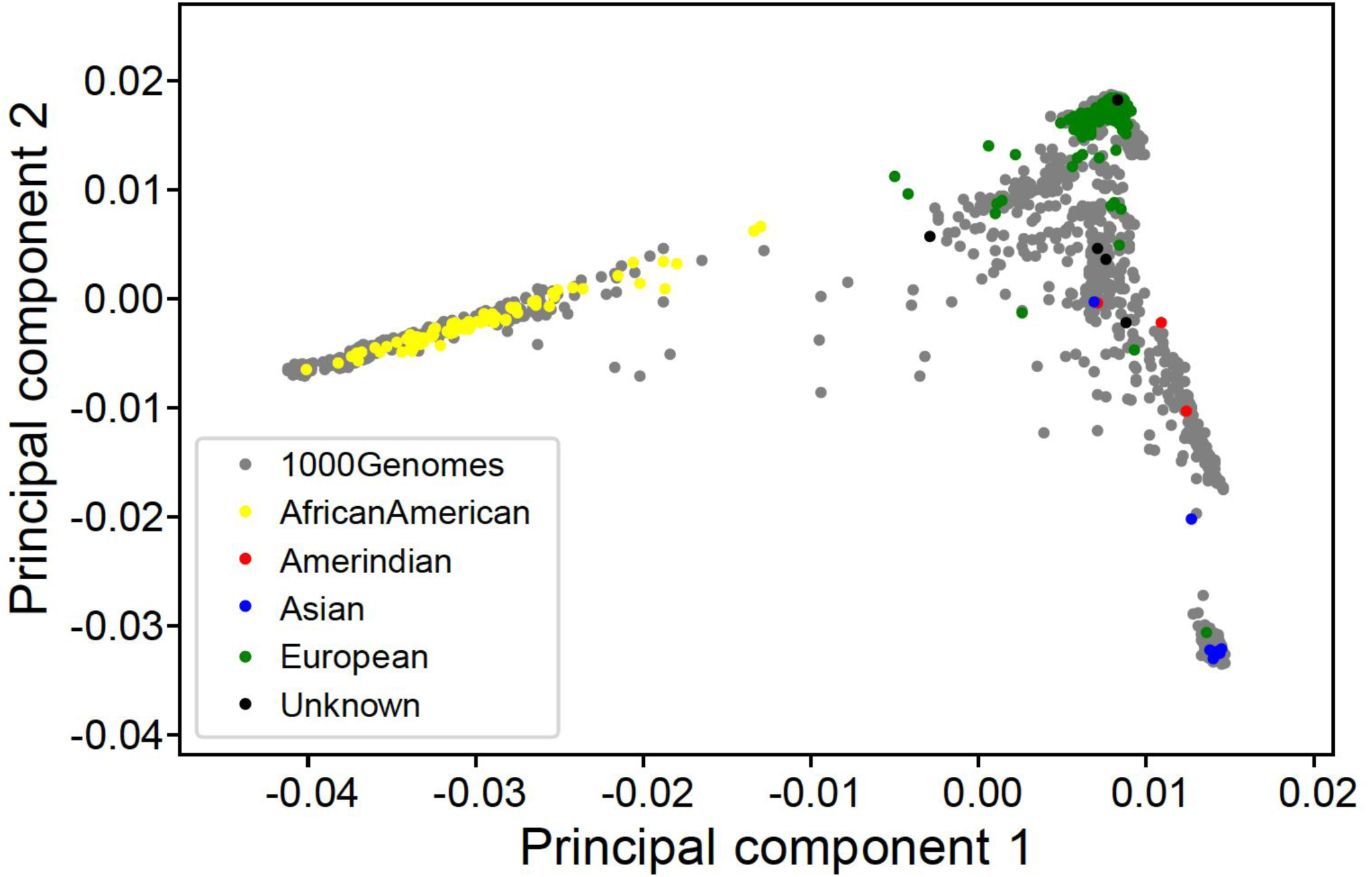
Analysis of GTEx population structure. Principal component analysis was performed using SNP genotypes from the GTEx and 1000 Genomes cohorts. Samples from the 1000 Genomes project are shown in gray and GTEx samples are shown as colored dots based on ethnicity provided for each sample (yellow=African American; red=Amerindian; blue=Asian; green=European, black=Unknown). Related to Fig. 1.

**Supplementary Figure 2:**
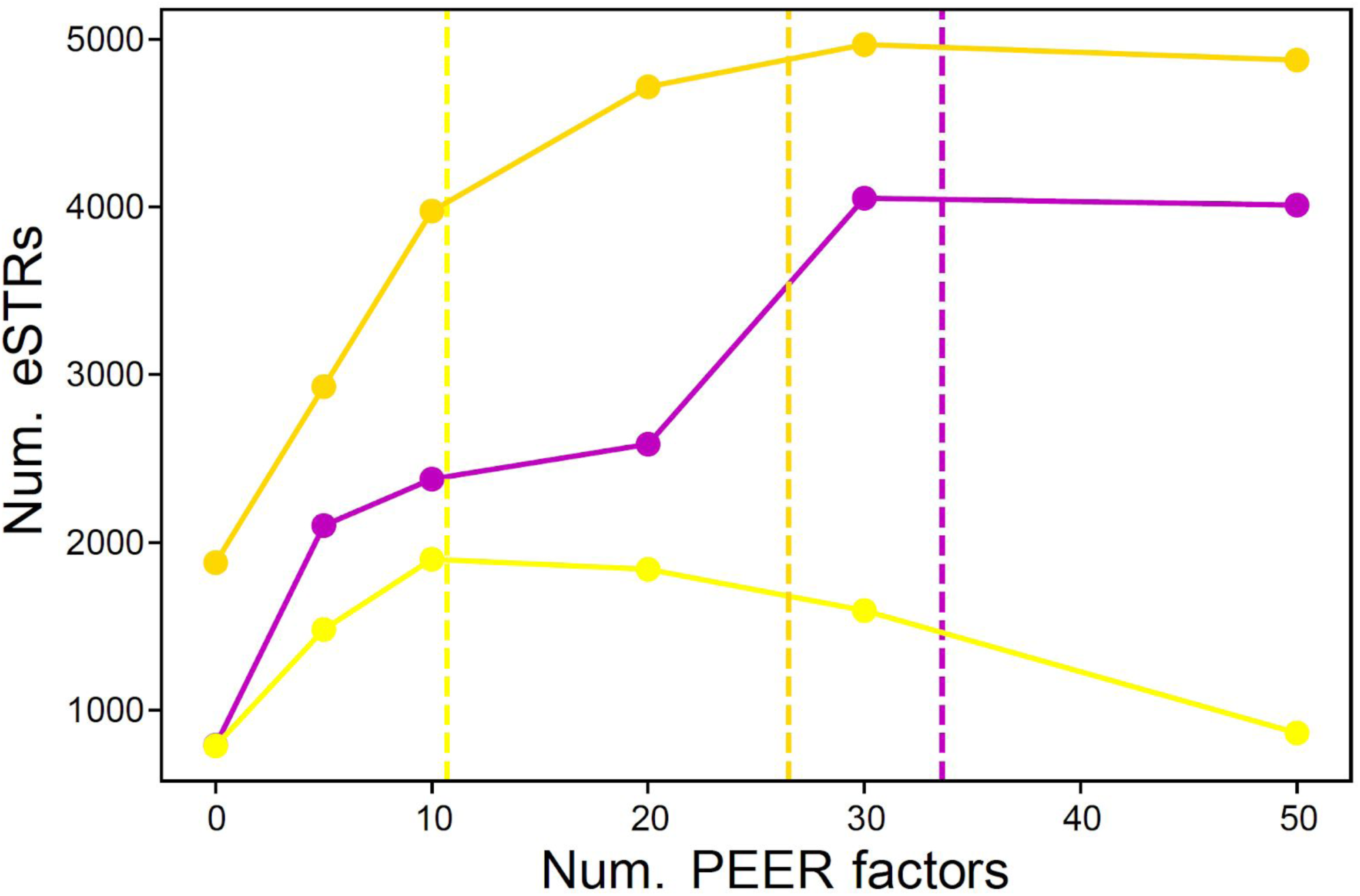
Effect of varying numbers of PEER factors on power to detect eSTRs. eSTRs (gene-level FDR 10%) were computed for each tissue after adjusting for a number of PEER factors ranging from 0 to 50. The x-axis shows the number of PEER factors adjusted for. The y-axis shows the number of significant eSTRs. Dashed vertical lines show the number of PEER factors equal to N/10, where N is the number of samples analyzed for each tissue. Purple=whole blood, yellow=Brain-Cerebellum, gold=Nerve-Tibial. Related to Fig. 1.

**Supplementary Figure 3:**
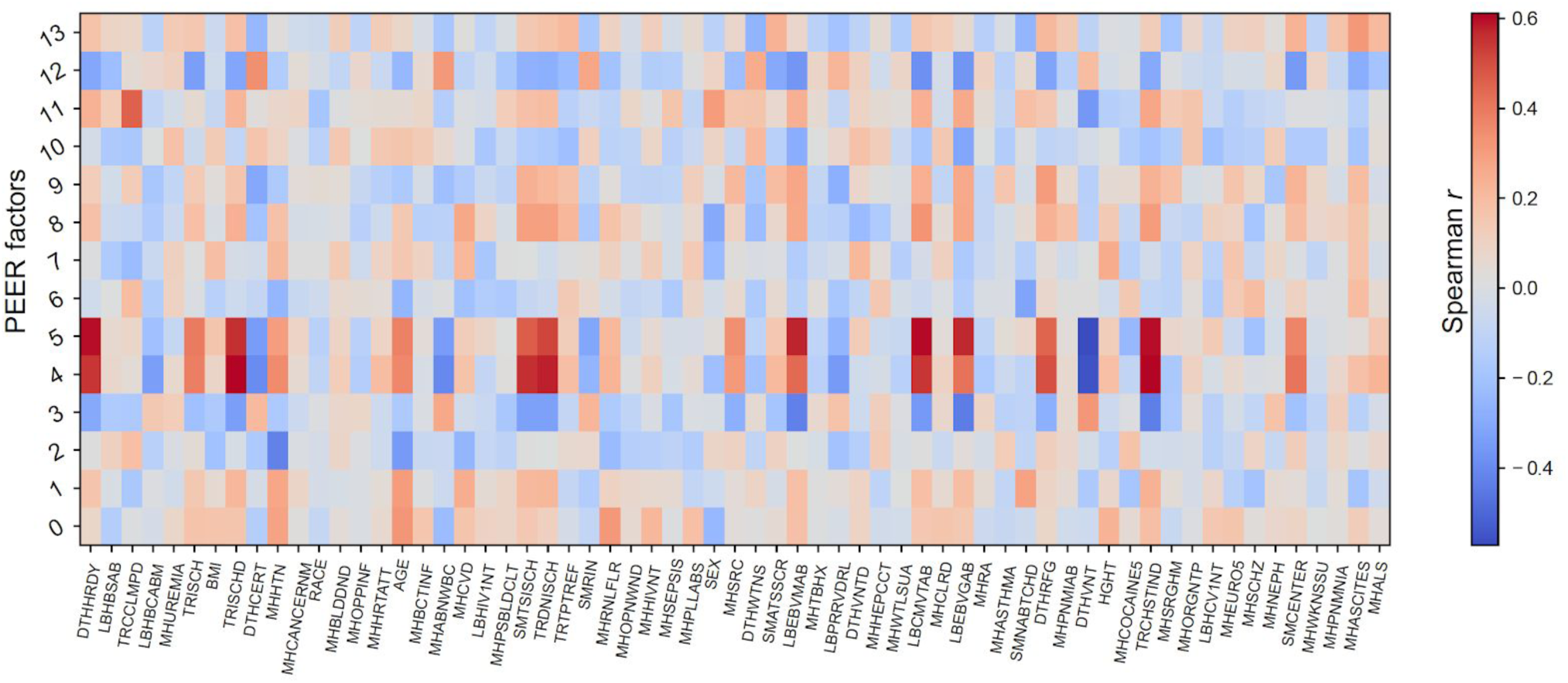
Correlation of sample metadata with PEER factors. Each cell in the matrix shows the Spearman correlation of each PEER factor with data processing covariates. The x-axis represents each variable as defined for the GTEx cohort in dbGaP study phs000424.v7.p2. For example, covariates most strongly associated with PEER factors included DTHHRDY (Hardy scale for death classification) and TRISCHD (ischemic time). The y-axis represents factors obtained from PEER analysis of gene expression from Adipose-subcutaneous tissue. Similar correlations were observed for other tissues. Related to Fig. 1.

**Supplementary Figure 4:**
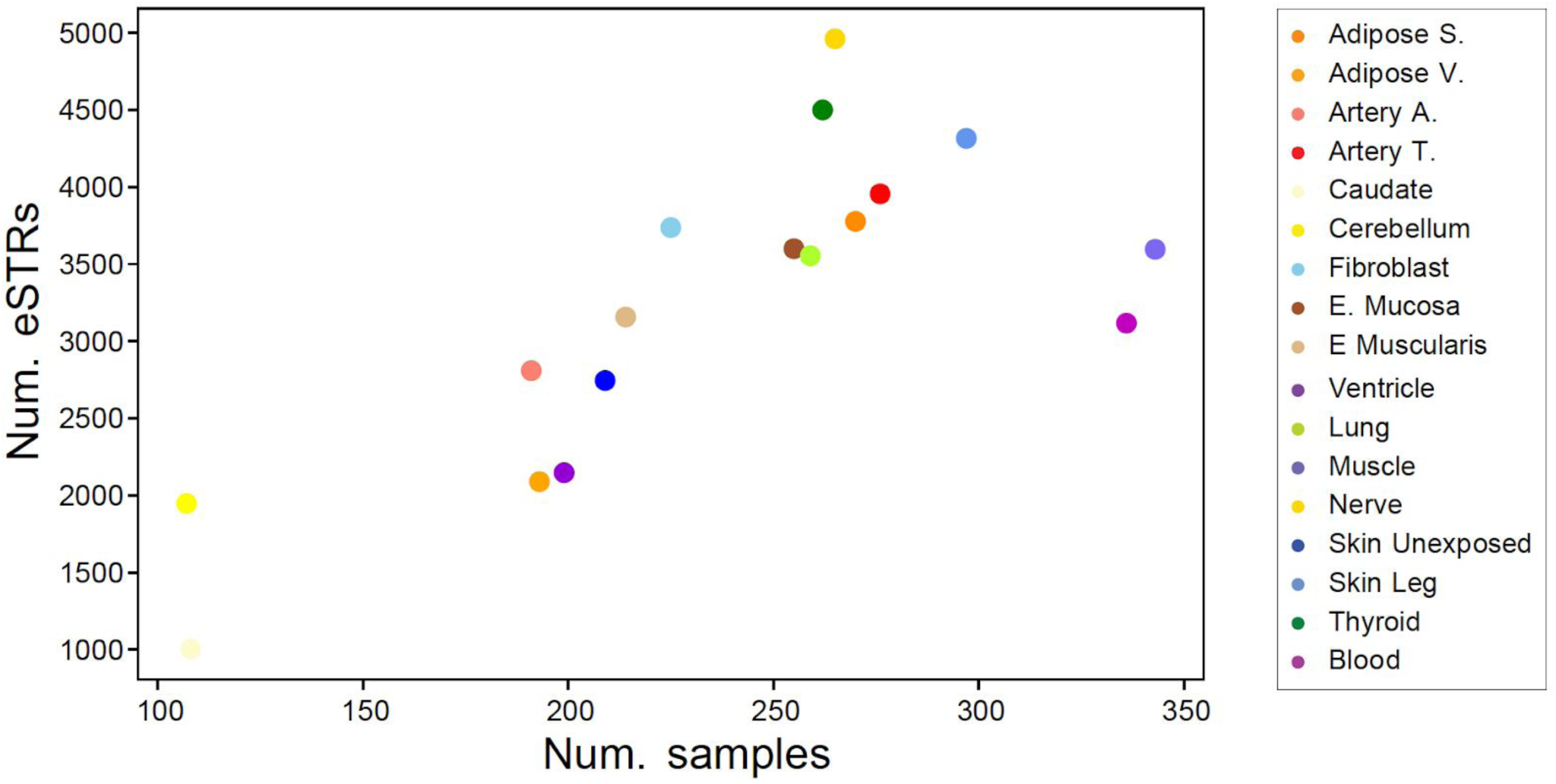
Relationship between sample size and number of eSTRs detected. The x-axis shows the number of samples per tissue. The y-axis shows the number of eSTRs (gene-level FDR<10%) detected in each tissue. Each dot represents a single tissue, using the same colors as shown in Fig. 1 in the main text (box on the right). Related to Fig. 1.

**Supplementary Figure 5:**
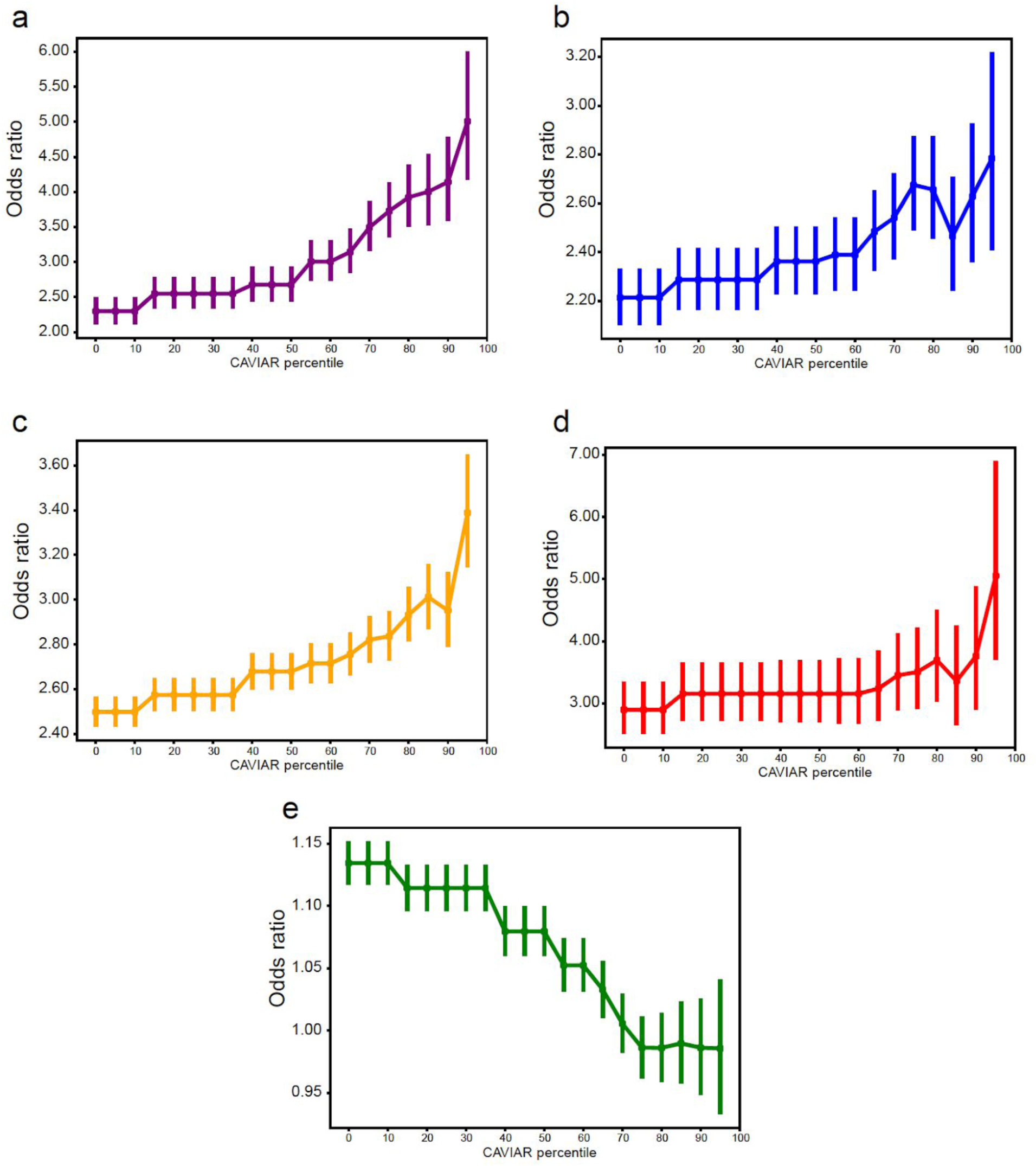
Enrichment of genomic annotations as a function of CAVIAR threshold. The x-axis represents CAVIAR threshold in terms of the percentile across all eSTRs. The y-axis represents the odds ratio for enrichment in eSTRs above each percentile threshold in each of these categories: **a.** 5’UTRs (purple); **b.** 3’UTRs (blue); **c.** promoters (orange; within 3kb of a transcription start site); **d.** Coding regions (red) and **e.** Intergenic regions (green). Related to Fig. 1.

**Supplementary Figure 6:**
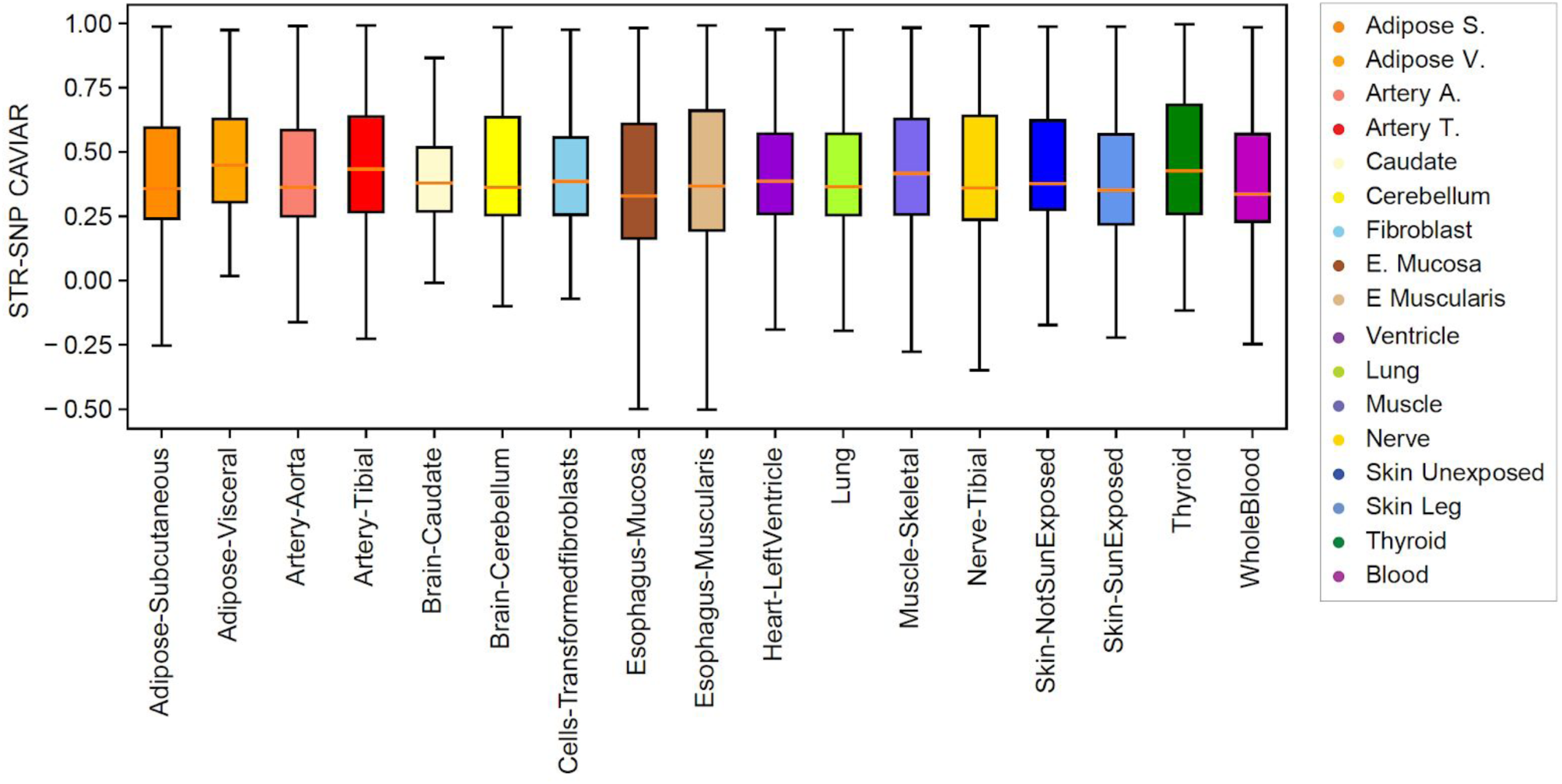
Difference in CAVIAR score between the top eSTR and top eSNP for each gene. For each tissue, the boxplot shows the distribution of differences between the CAVIAR posterior score for the best STR vs. the best SNP for each gene. Data is only shown for genes with FM-eSTRs. The colors of each box correspond to the different tissues (see legend on the right and the same as in Fig. 1b). Boxplots as in Fig. 1c. Outlier points are not shown. Related to Fig. 1.

**Supplementary Figure 7:**
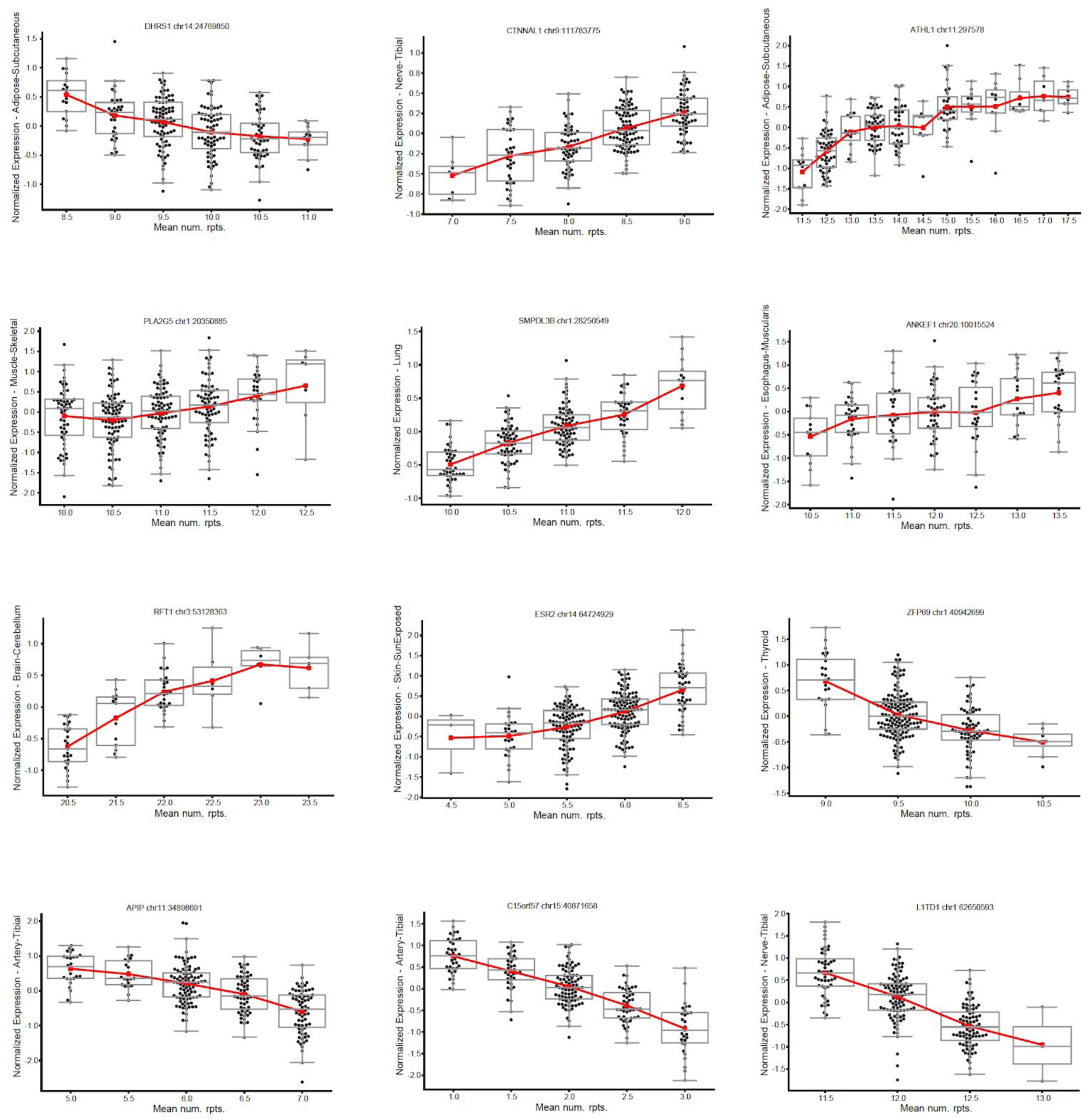
Example multi-allelic FM-eSTRs. For each plot, the x-axis represents the mean number of repeats in each individual and the y-axis represents normalized expression in the tissue for which the eSTR was most significant. Boxplots summarize the distribution of expression values for each genotype. Boxplots as in Fig. 1c.The red line shows the mean expression for each x-axis value. Related to Fig. 1.

**Supplementary Figure 8:**
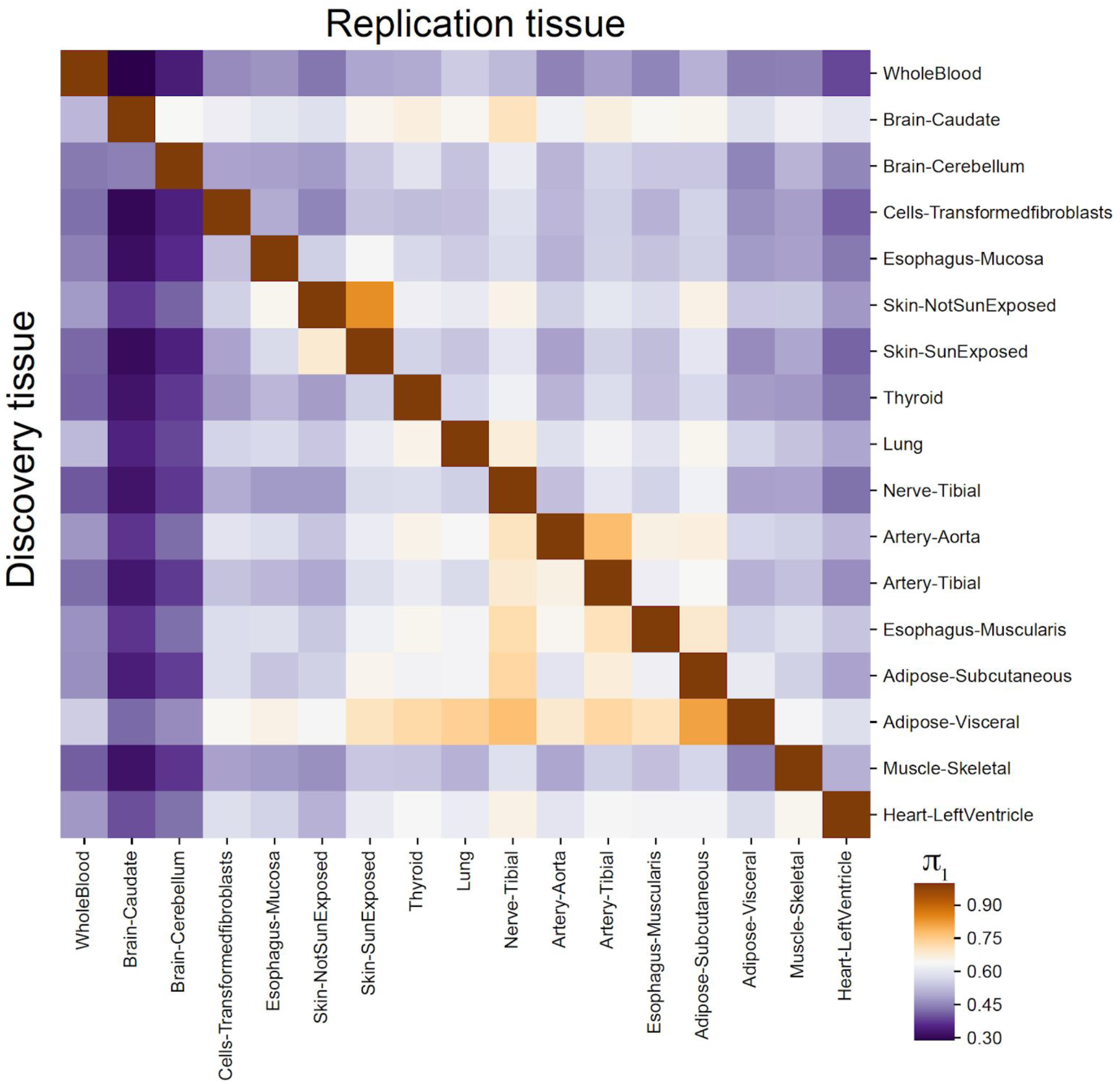
Pairwise sharing of effect sizes across tissues. For each discovery tissue (rows), all eSTRs with gene-level FDR<10% were tested for association in each other (replication) tissue (columns). The value in each cell gives the percent of eSTRs that were replicated with p<0.05 (π_1_). Related to Fig. 1.

**Supplementary Figure 9:**
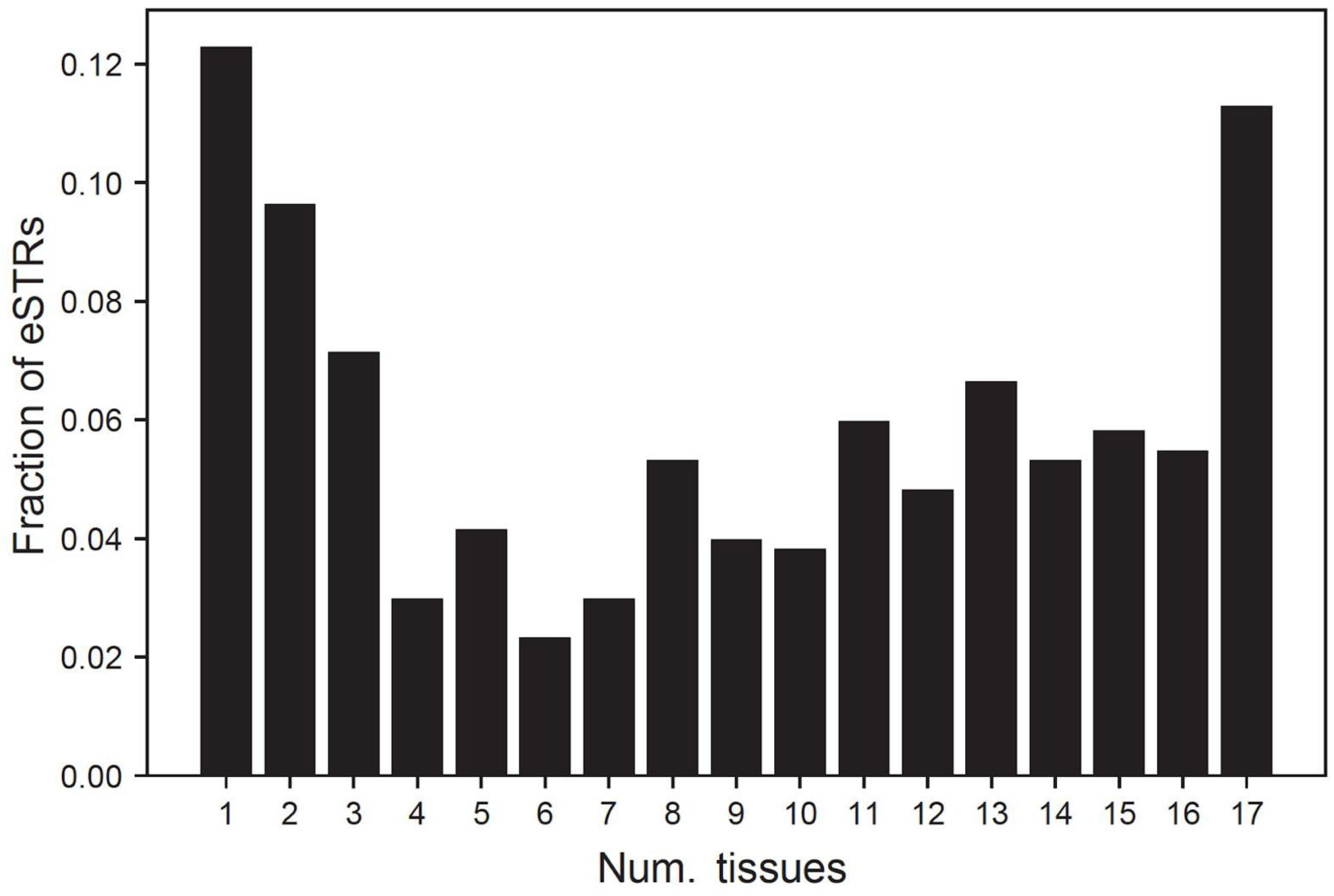
Sharing of eSTRs across tissues. The x-axis represents the number of tissues that share a given eSTR (absolute value of mashR Z-score >4). The y-axis represents the number of eSTRs shared across a given number of tissues. Related to Fig. 1.

**Supplementary Figure 10:**
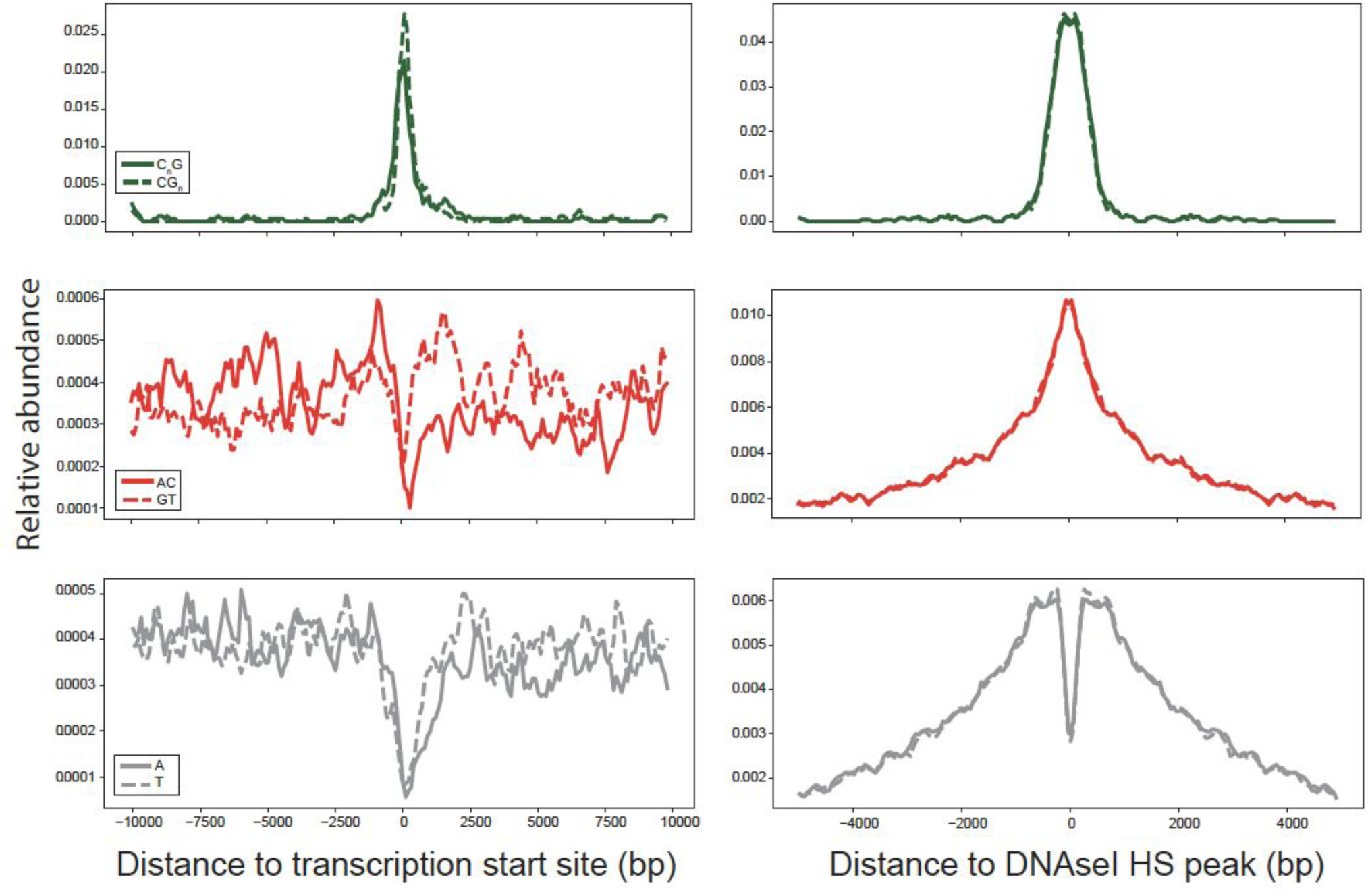
Localization of all STRs around putative regulatory regions. Left and right plots show localization around transcription start sites and DNAseI HS clusters, respectively. The y-axis denotes the relative number of STRs of each type in each bin. For promoters, the x-axis is divided into 100bp bins. For DNAseI HS sites, the x-axis is divided into 50bp bins. In each plot, values were smoothed by taking a sliding average of each four consecutive bins. Only STR-gene pairs included in our analysis are considered. Each plot compares localization of the two possible sequences of a given repeat unit on the coding strand. *i.e.* top plots compare repeat units of the form C_n_G vs. their reverse complement on the opposite strand, middle plots compare AC vs. GT repeats, and bottom plots compare A vs. T repeats. The strand of each STR was determined based on the coding strand of each target gene. Related to Fig. 2.

**Supplementary Figure 11:**
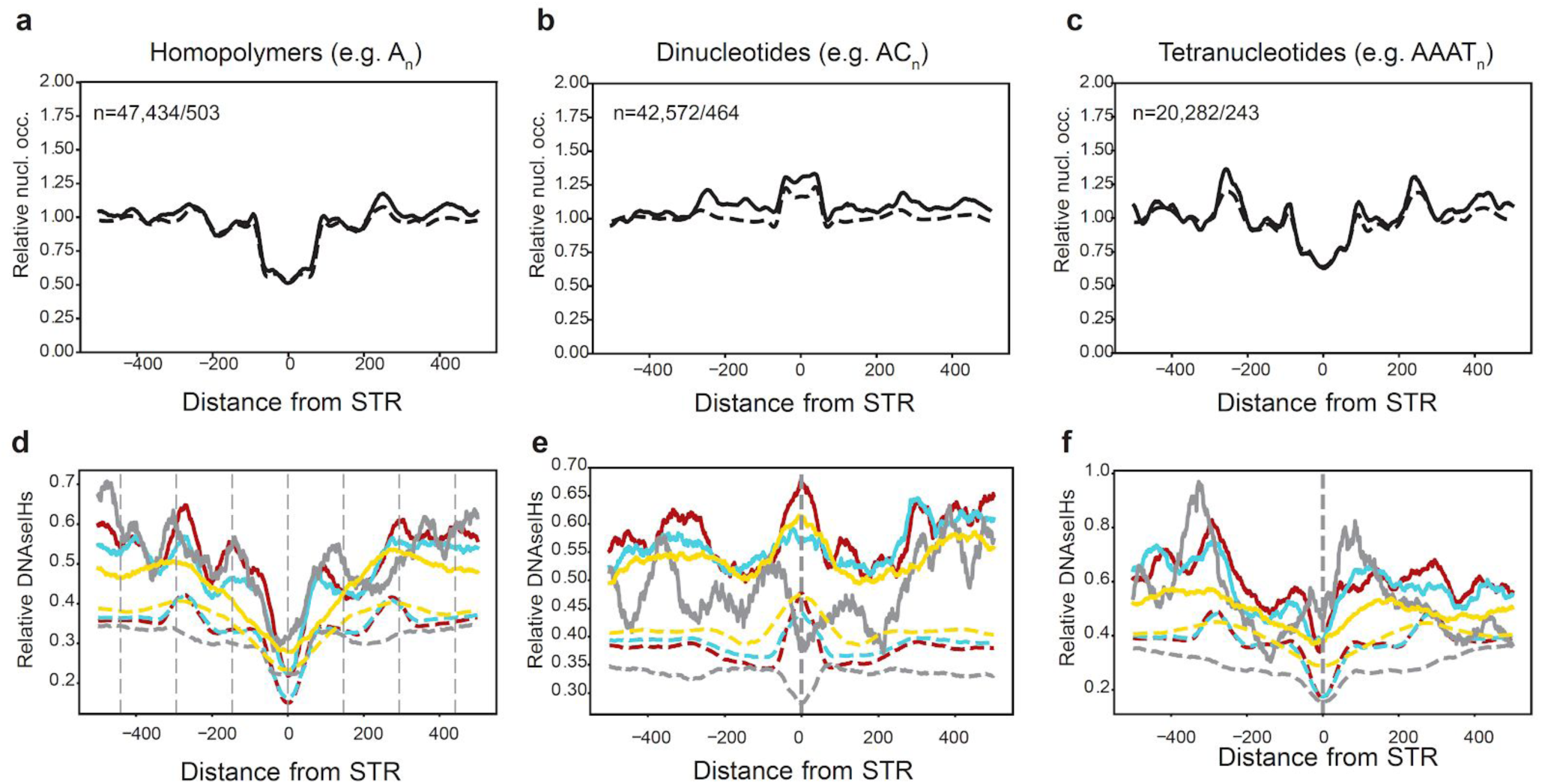
Nucleosome occupancy and DNAseI hypersensitivity show distinct patterns around eSTRs a-c. Nucleosome density around STRs with different repeat unit lengths. Nucleosome density in GM12878 in 5bp windows is averaged across all STRs analyzed (dashed) and FM-eSTRs (solid) relative to the center of the STR. **b. DNAseI HS density around STRs with different repeat unit lengths.** The number of DNAseI HS reads in GM12878 (gray), fat (red), tibial nerve (yellow), and skin (cyan) is averaged across all STRs in each category. Solid lines show FM-eSTRs. Dashed lines show all STRs. Left=homopolymers, middle=dinucleotides, right=tetranucleotides. Other repeat unit lengths were excluded since they have low numbers of FM-eSTRs (see Fig. 5a). Dashed vertical lines in **(d)** show the STR position +/- 147bp. Related to Fig. 2.

**Supplementary Figure 12:**
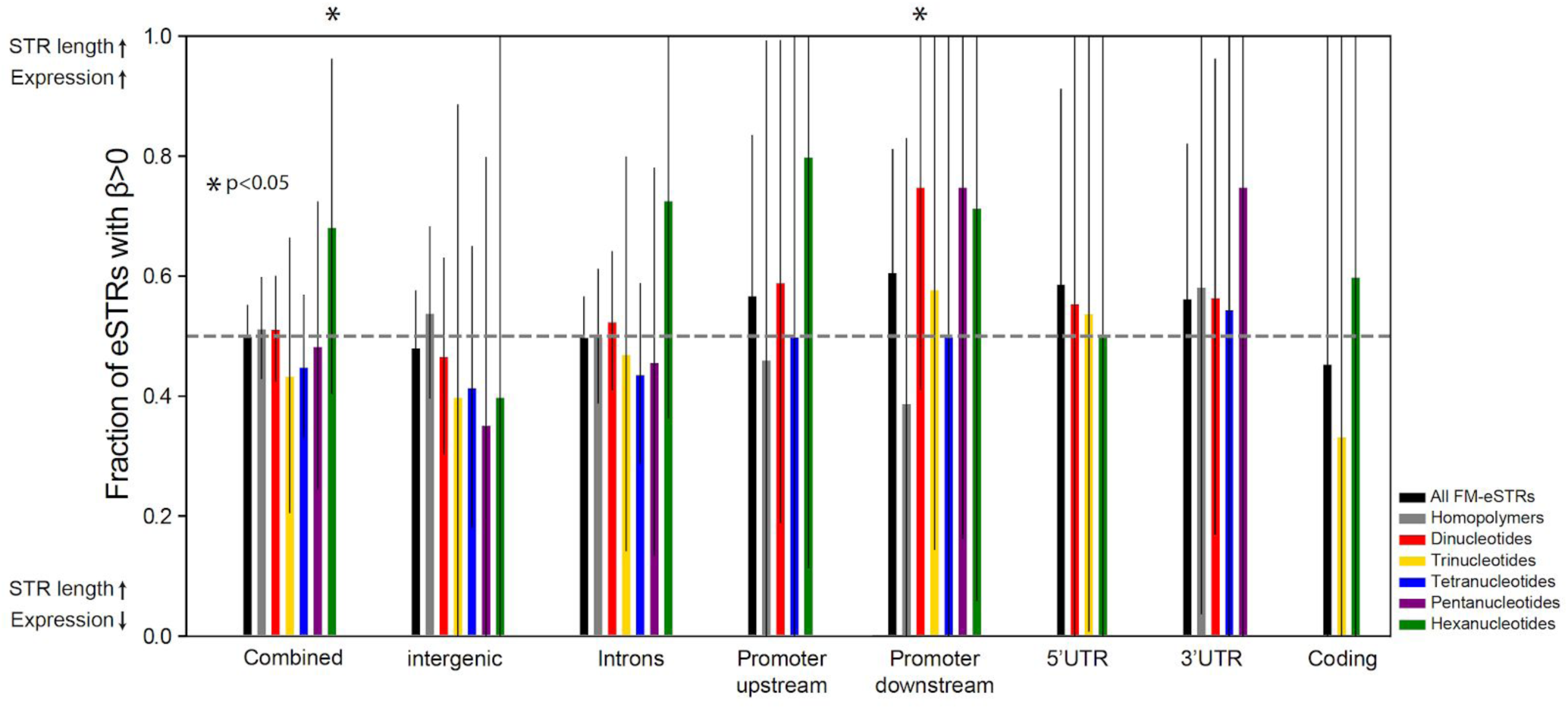
Bias in the direction of eSTR effect sizes. The y-axis shows the percentage of FM-eSTRs in each category with positive effect sizes, meaning a positive correlation between STR length and expression. Colored bars represent different repeat unit lengths (black=all FM-eSTRs; gray=homopolymers; red=dinucleotides; gold=trinucleotides; blue=tetranucleotides; purple=pentanucleotides; green=hexanucleotides). Error bars show 95% confidence intervals. Asterisks denote categories that are nominally significant (binomial two-sided p<0.05) for having significantly more or less positive effect sizes than expected by chance (50%). No category was significant after accounting for multiple hypothesis testing. Related to Fig. 2.

**Supplementary Figure 13:**
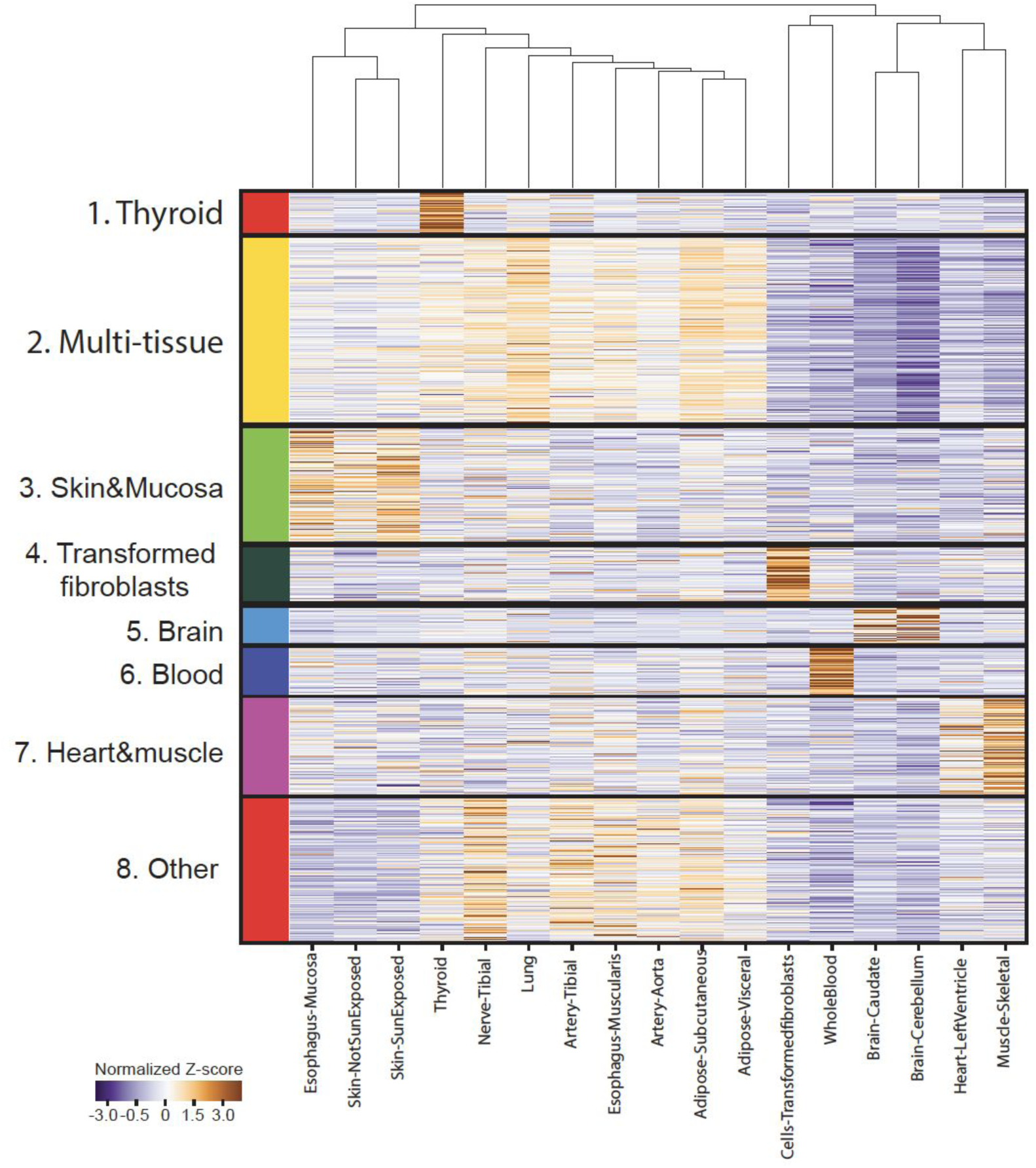
Characterization of tissue-specific FM-eSTRs. FM-eSTRs were clustered by absolute Z-scores computed by mashR using K-means (**Methods**). The heatmap shows absolute values of Z-scores in each tissue, Z-normalized by row. (Number of genes in each cluster: Cluster 1=22, Cluster 2=497, Cluster 3=253; Cluster 4=122, Cluster 5=90, Cluster 6=126, Cluster 7=220, Cluster 8=336). Related to Fig. 2.

**Supplementary Figure 14:**
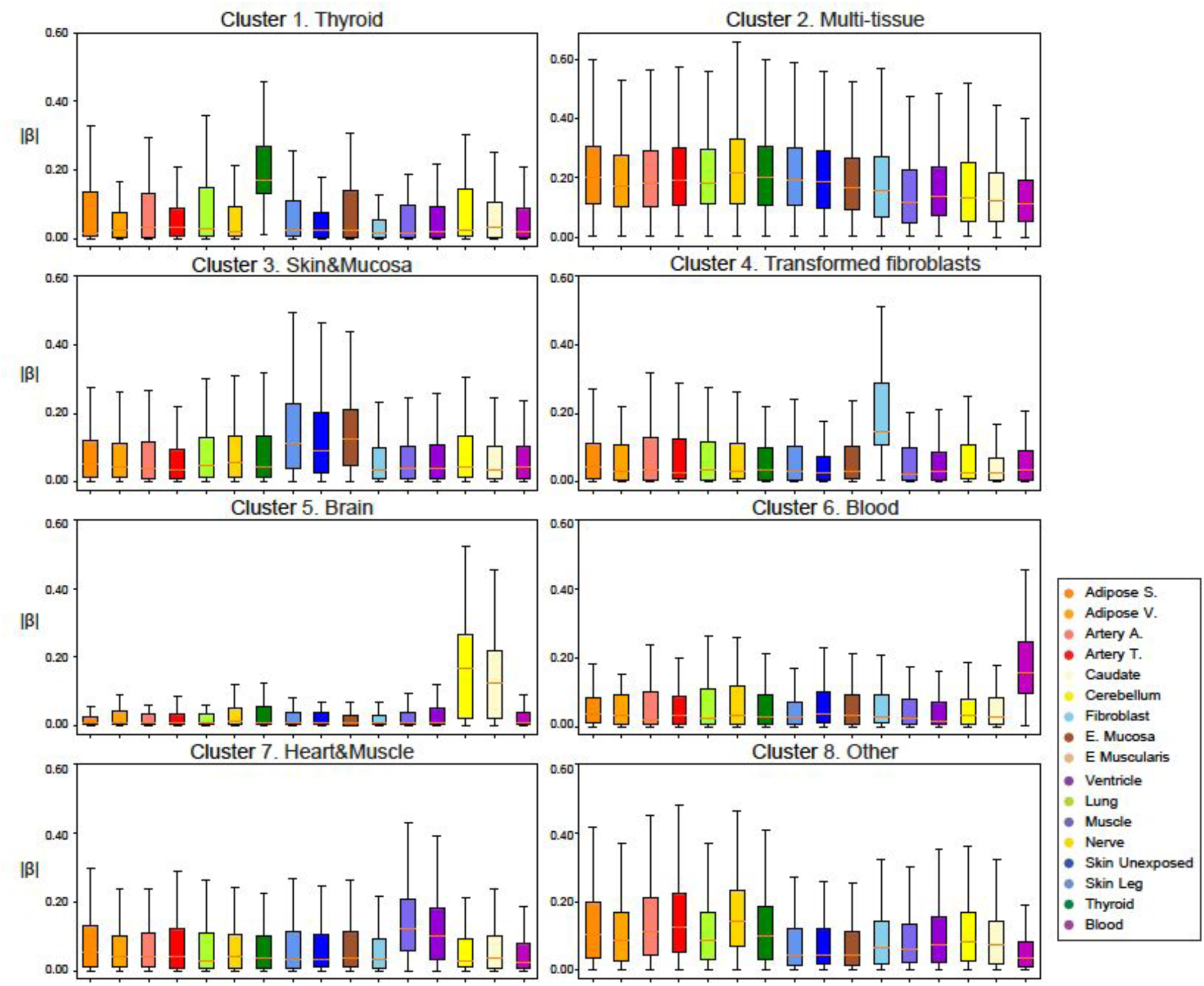
Characterization of tissue-specific eSTRs. Each panel shows the distribution of the absolute value of posterior effect sizes computed by mashR in each tissue for the set of FM-eSTRs in each cluster (see Supp Fig. 13 above). Horizontal lines show median values, boxes span from the 25th percentile (Q1) to the 75th percentile (Q3). Whiskers extend to Q1-1.5*IQR (bottom) and Q3+1.5*IQR (top), where IQR gives the interquartile range (Q3-Q1). The red line shows the mean expression for each x-axis value. Related to Fig. 2.

**Supplementary Figure 15:**
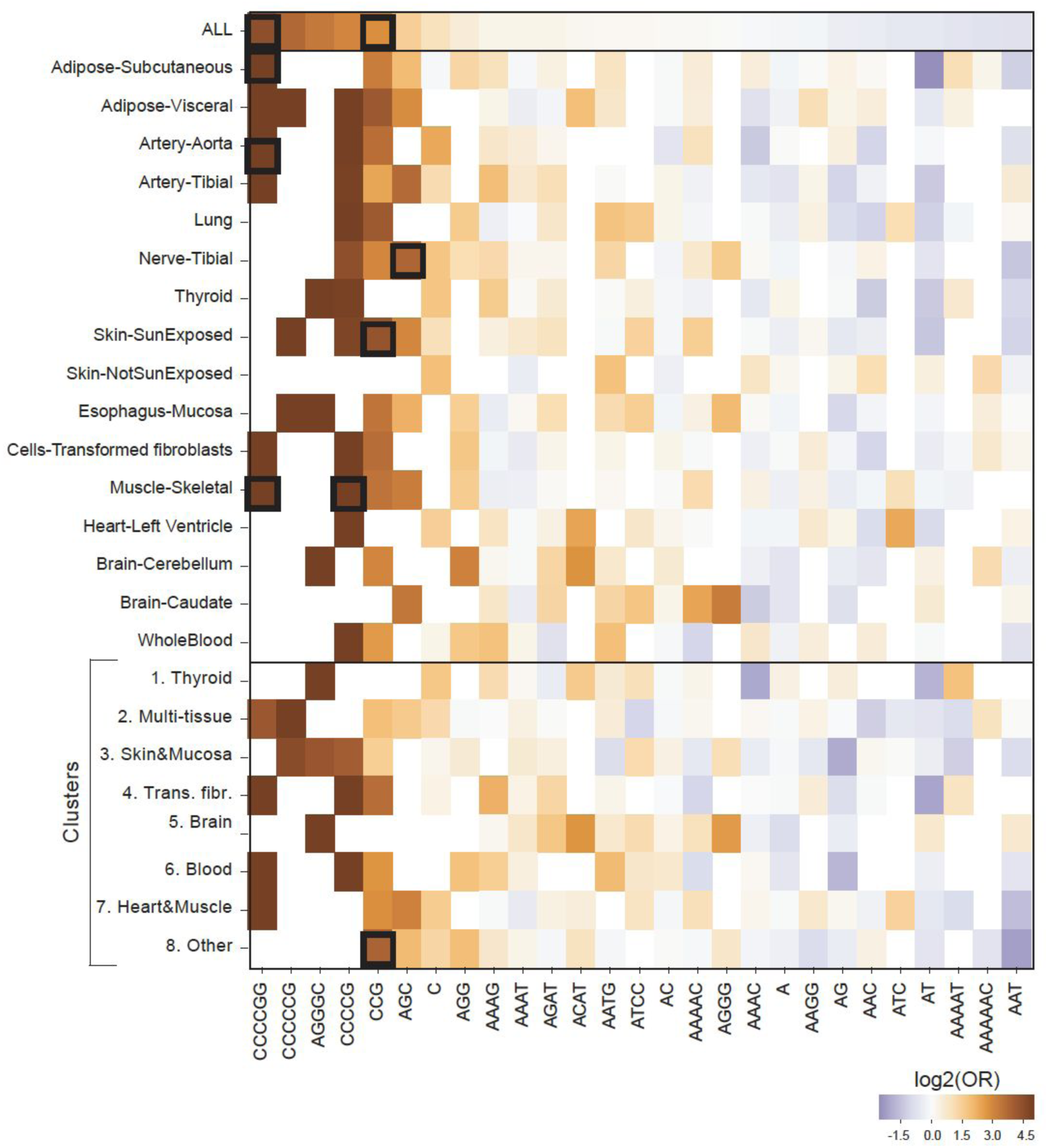
eSTR repeat unit enrichment. We evaluated repeat unit enrichment in multiple FM-eSTR groups: all FM-eSTRs combined across tissues (similar to Fig. 2e), FM-eSTRs identified per-tissue, and FM-eSTRs belonging to each cluster (see Supplementary Fig. 13). For each group of FM-eSTRs, the heatmap shows the log2 of the odds ratio computed using a Fisher’s Exact test (scipy.stats.fisher_exact). Columns are sorted from highest to lowest enrichment in all FM-eSTRs. Bold boxes indicate enrichments statistically significant (adjusted p<0.05, adjusted separately per row for the number of motifs tested). Related to Fig. 2.

**Supplementary Figure 16:**
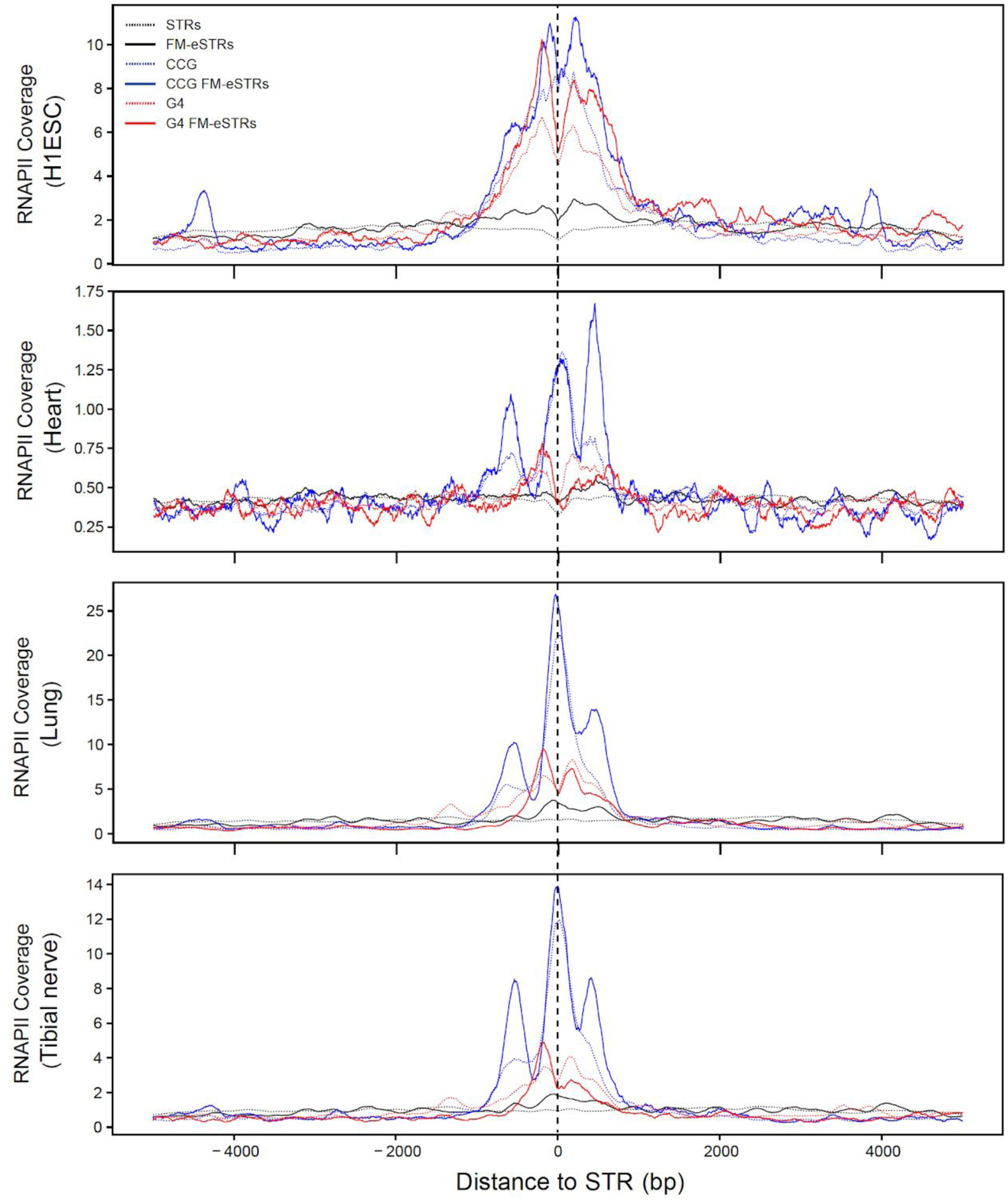
Density of RNAPII localization around STRs. The y-axis shows the average number of ChIP-seq reads for RNA Polymerase II in 5bp bins centered at STRs within 5KB of TSSs. Black lines denote all STRs, blue lines denote CCG STRs, and red lines denote STRs matching the canonical G4 motif. Dashed lines represent all STRs of each class and solid lines represent FM-eSTRs. Plots show read counts in different cell types. From top to bottom: human embryonic stem cells, heart, lung, and tibial nerve. Related to Fig. 3.

**Supplementary Figure 17:**
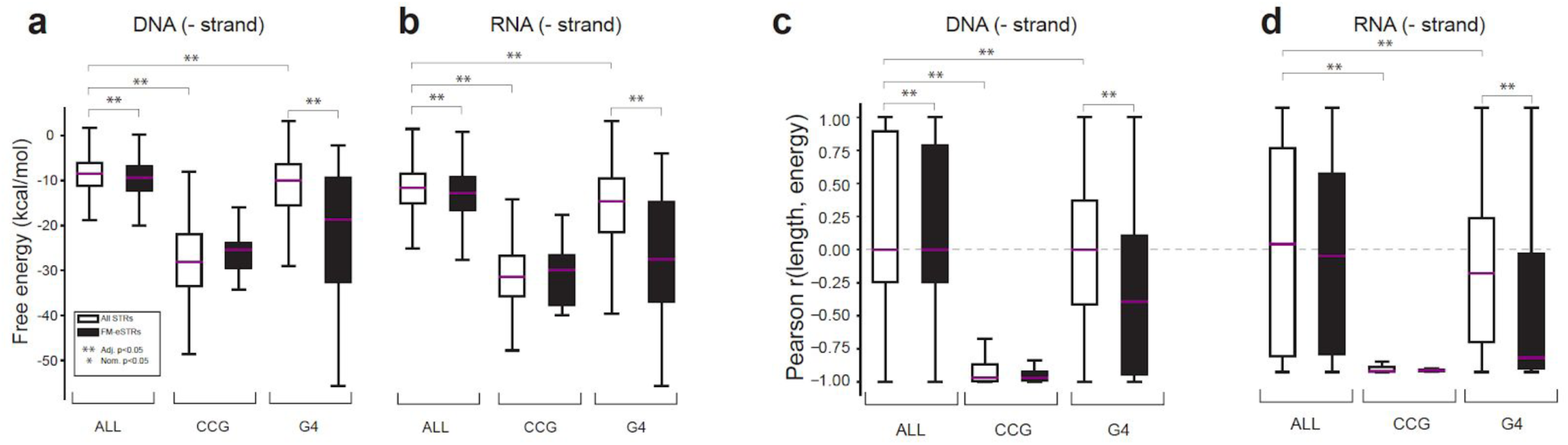
GC-rich eSTRs modulate DNA or RNA secondary structure. (**a-b) Free energy of STR regions.** Boxplots denote the distribution of free energy for each STR +/- 50bp of context sequence, computed as the average across all alleles at each STR. **(a)** and **(b)** show results computed using the non-template strand for DNA and RNA respectively. (**c-d) Pearson correlation between STR length and free energy.** Correlations were computed separately for each STR, and plots show the distribution of correlation coefficients across all STRs. The dashed horizontal line denotes 0 correlation as expected by chance. **(c)** and **(d)** show results computed using the non-template strand for DNA and RNA respectively. Nominally significant (Mann Whitney one-sided p<0.05) differences between distributions are denoted with a single asterisk. Differences significant after controlling for multiple hypothesis correction are denoted with double asterisks. For each category (free energy and Pearson correlation), we used a Bonferroni correction to control for 20 total comparisons: comparing all vs. FM-eSTRs separately in each category, comparing CCG vs. all STRs, and comparing G4 vs. all STRs, in four conditions (DNA +/- and RNA +/-). Related to Fig. 3.

**Supplementary Figure 18:**
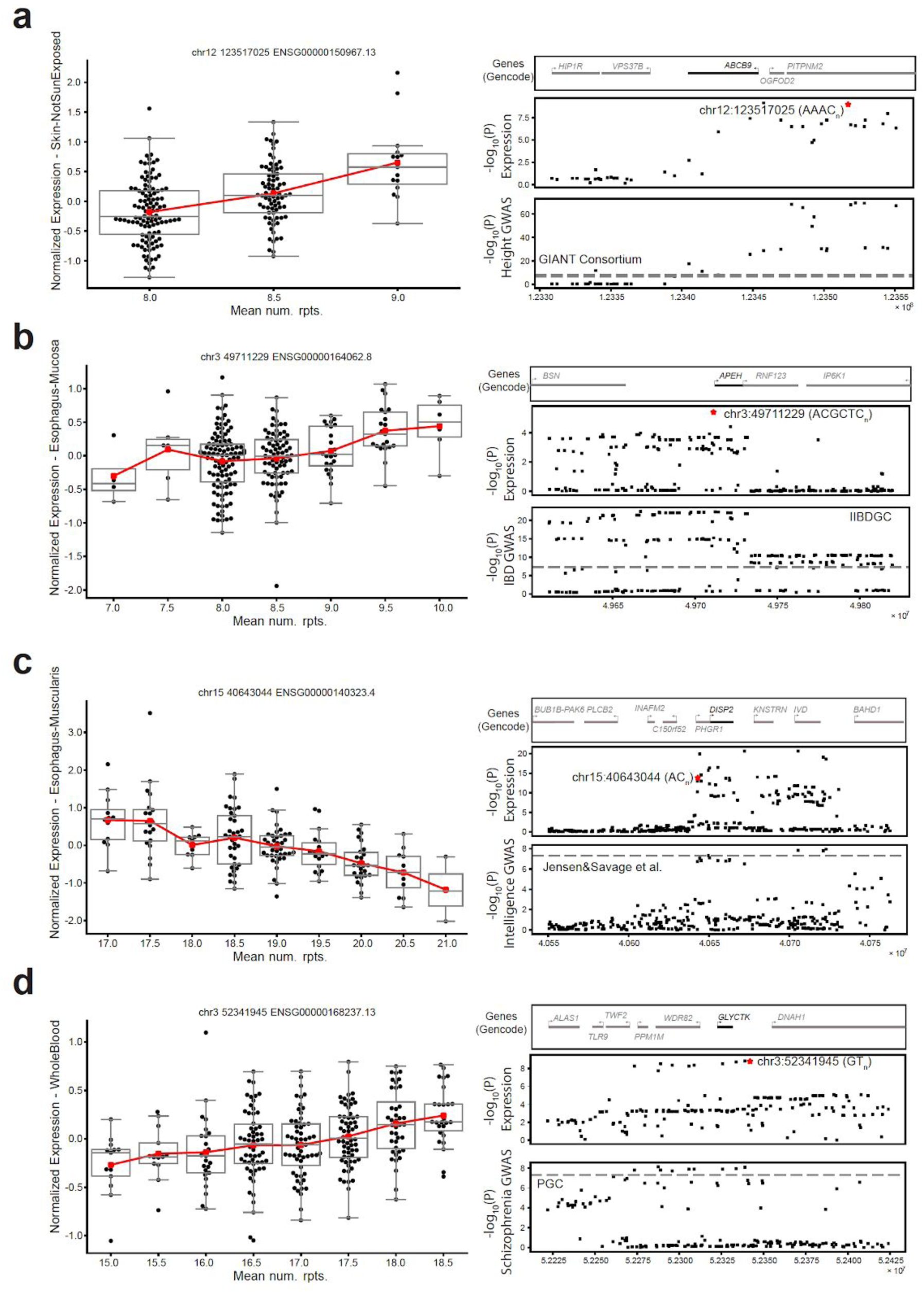
Example GWAS signals co-localized with FM-eSTRs. Left: For each plot, the x-axis represents the mean number of repeats in each individual and the y-axis represents normalized expression in the tissue with the most significant eSTR signal at each locus. Boxplots summarize the distribution of expression values for each genotype. Box plots as in Fig. 1c. The red line shows the mean expression for each x-axis value. Right: Top panels give genes in each region. The target gene for the eQTL associations is shown in black. Middle panels give the -log_10_ p-values of association of the effect-size between each SNP (black points) and the expression of the target gene. The FM-eSTR is denoted by a red star. Bottom panels give the -log_10_ p-values of association between each SNP and the trait based on published GWAS summary statistics. Dashed gray horizontal lines give the genome-wide significance threshold of 5E-8. Related to Fig. 4.

**Supplementary Figure 19:**
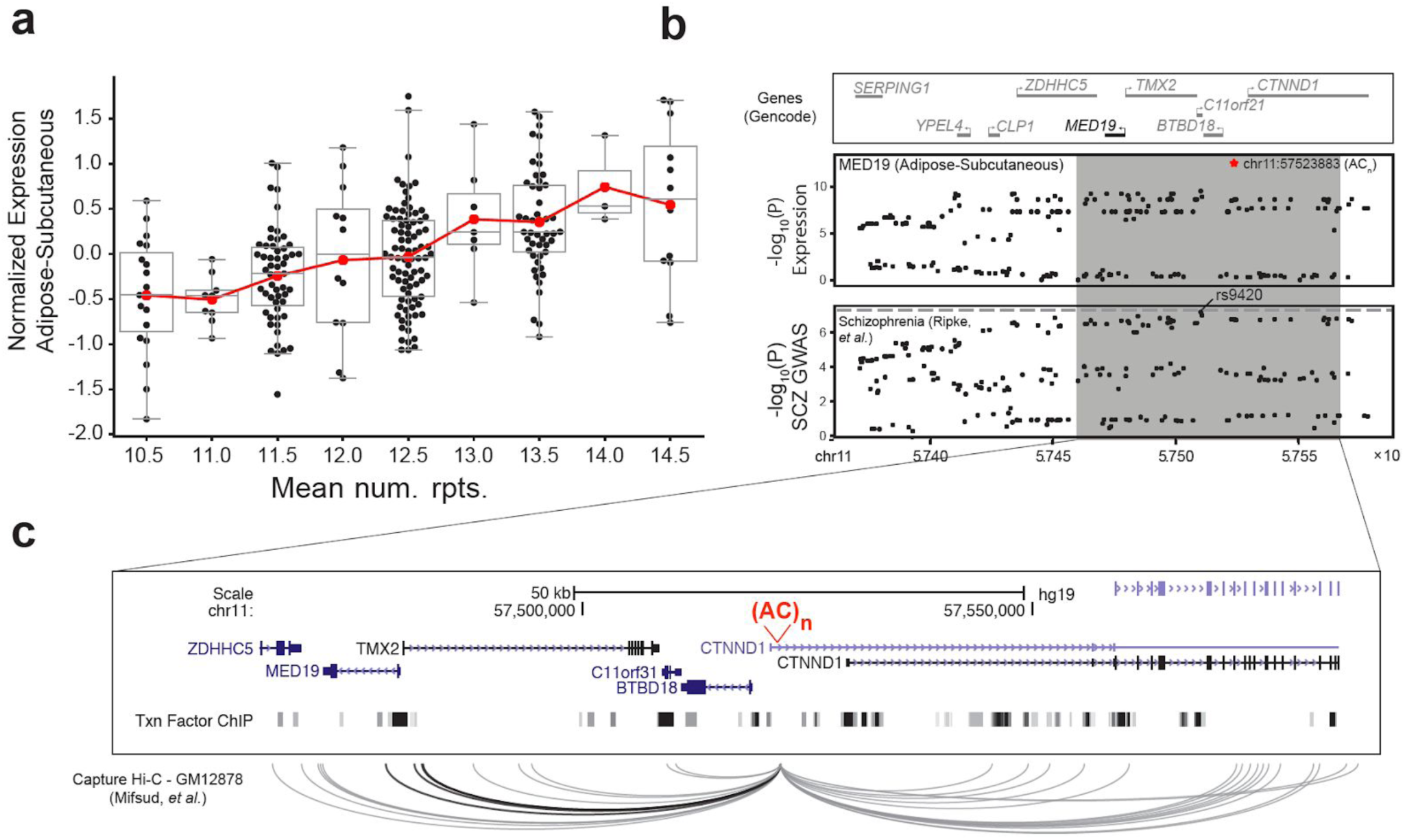
Example GWAS signal for schizophrenia potentially driven by an eSTR for *MED19*. **a. eSTR association for *MED19*.** The x-axis shows STR genotypes at an AC repeat (chr11:57523883) as the mean number of repeats in each individual and the y-axis shows normalized *MED19* expression in subcutaneous adipose. Each point represents a single individual. Red lines show the mean expression for each x-axis value. Boxplots as in Fig. 1c. **b. Summary statistics for *MED19* expression and schizophrenia.** The top panel shows genes in the region around *MED19*. The middle panel shows the -log_10_ p-values of association between each variant and *MED19* expression in subcutaneous adipose tissue in the GTEx cohort. The FM-eSTR is denoted by a red star. The bottom panel shows the -log_10_ p-values of association for each variant with schizophrenia from Ripke, *et al.*. The dashed gray horizontal line shows genome-wide significance threshold of 5E-8. **c. Detailed view of the *MED19* locus.** A UCSC genome browser screenshot is shown for the region in the gray box in **(b)**. The FM-eSTR is shown in red [(AC)n]. The bottom track shows transcription factor (TF) and chromatin regulator binding sites profiled by ENCODE. The bottom panel shows long-range interactions reported by Mifsud, *et al.* using Capture Hi-C on GM12878. Interactions shown in black include *MED19*. Interactions to loci outside of the window depicted are not shown. Related to Fig. 4. See file eSTRGtex_SuppTables.xlsx for Supplementary Tables 1-8.

