## Supplemental Dataset 2 for "Multi-tissue analysis reveals short tandem repeats as ubiquitous regulators of gene expression and complex traits": README

Each file contains eSTR statistics for each tissue analyzed. The columns are described below:

1. chrom: chromosome (hg19)
2. gene: ENSEMBL gene ID
3. str.start: start position of the STR (hg19)
4. mashr.beta: posterior estimate of effect size from mashR
5. mashr.beta.se: standard error of effect size from mashR
6. mashr.top.str: indicates if this is the top STR for the gene based on mashR Z-scores
7. mashr.significant: indicates whether the mashR Z-score has absolute value >= 3
8. str.end: end position of the STR (hg19)
9. str.motif.forward: repeat unit of the STR on the forward reference strand
10. str.motif.reverse: repeat unit of the STR on the reverse reference strand
11. linreg.beta: Effect size of the eSTR estimated by linear regression
12. linreg.beta.se: Standard error of the effect size of the eSTR estimated by linear regression
13. linreg.pval: Linear regression p-value for the eSTR
14. linreg.n.miss: Number of samples not included in the linear regression analysis
15. linreg.absZ: Absolute value of the Z-score from linear regression
16. linreg.top.pval: The best p-value for all STRs tested for this gene
17. linreg.top.absZ: The best Z-score (absolute value) for all STRs tested for this gene
18. num.strs: Number of STRs analyzed for this gene in the linear regression
19. linreg.qval: Gene-level false discovery rate for the top eSTR for each gene
20. linreg.top.str: Indicates whether the STR was the top eSTR for the gene based on linear regression
21. linreg.significant: Indicates whether the STR was significant based on linear regression (gene-level FDR of 10%)
22. mashr.top.snp: The top SNP from mashR results based on Z-score
23. anova.pval: P-value of the ANOVA analysis comparing (Y~SNP) vs. (Y~SNP+STR)
24. anova.qval: Adjusted p-value of the ANOVA analysis
25. anova.best.snp: The ID of the SNP included in the ANOVA analysis
26. caviar.str.score: CAVIAR score for the STR
27. caviar.str.rank: Rank of the STR's CAVIAR score compared to all other variants analyzed by CAVIAR
28. caviar.topsnp: Top SNP based on CAVIAR analysis
29. caviar.topsnp.score: CAVIAR score of the top SNP
30. caviar.nsnps: Number of SNPs included in the CAVIAR analysis
31. gene.name: Name of the gene
32. gene.strand: Strand of the gene
